# Proteome constraints reveal targets for improving microbial fitness in nutrient-rich environments

**DOI:** 10.1101/2020.10.15.340554

**Authors:** Yu Chen, Eunice van Pelt-KleinJan, Berdien van Olst, Sieze Douwenga, Sjef Boeren, Herwig Bachmann, Douwe Molenaar, Jens Nielsen, Bas Teusink

**Author notes:** These authors contributed equally: Yu Chen, Eunice van Pelt-KleinJan. These authors jointly supervised this work: Jens Nielsen, Bas Teusink.

## Abstract

Cells adapt to different conditions via gene expression that tunes metabolism and stress resistance for maximal fitness. Constraints on cellular proteome may limit such expression strategies and introduce trade-offs^1^; Resource allocation under proteome constraints has emerged as a powerful paradigm to explain regulatory strategies in bacteria^2^. It is unclear, however, to what extent these constraints can predict evolutionary changes, especially for microorganisms that evolved under nutrient-rich conditions, i.e., multiple available nitrogen sources, such as the lactic acid bacterium *Lactococcus lactis*. Here we present an approach to identify preferred nutrients from integration of experimental data with a proteome-constrained genome-scale metabolic model of *L. lactis* (pcLactis), which explicitly accounts for gene expression processes and associated constraints. Using glucose-limited chemostat data^3^, we identified the uptake of glucose and arginine as dominant constraints, whose pathway proteins were indeed upregulated in evolved mutants. However, above a growth rate of 0.5 h^-1^, pcLactis suggests that available enzymes function at their maximum capacity, which allows an increase in growth rate only by altering gene expression to change metabolic fluxes, as was mainly observed for arginine metabolism. Thus, our integrative analysis of flux and proteomics data with a proteome-constrained model is able to identify and explain the constraints that form targets of regulation and fitness improvement in nutrient-rich growth environments.

---

The fitness of unicellular organisms is determined by adaptions to environmental conditions^1^, and is optimized by regulating metabolic processes that generally lead to higher growth rates^4^. Growth rates are finite, as the metabolic processes supporting growth are constrained through limits imposed by external conditions, e.g., nutrient availability, and internal factors that relate to cell morphology, enzyme kinetics and physicochemical properties such as solvent capacities. In particular, constraints on the allocation of the proteome, due to limited membrane area or intracellular volume, have aided to understand metabolic adaptions of microorganisms^5–7^, specifically, the overflow metabolism in *Escherichia coli*^8–10^ and the Crabtree effect in *Saccharomyces cerevisiae*^11–13^.

However, much less is known about metabolic adaptations in other organisms, and especially those cultivated under conditions with multiple available substrates, such as in nutrient-rich environments like the gut or food. It was previously shown that anaerobic overflow metabolism in the lactic acid bacterium *L. lactis* is not accompanied by changes in associated protein levels^3^, questioning the generality of the resource allocation paradigm. However, changes in gene expression were observed, in particular in amino acid metabolism, prompting us to revisit the cellular economics of *L. lactis*.

*L. lactis* is an important model lactic acid bacterium and work horse for the dairy industry - production of cheese in particular^14,15^. Many strains are auxotrophic for at least five amino acids^16,17^, and thus *L. lactis* strains are grown in nutrient-rich environments where amino acids are usually in excess. Many of these amino acids participate not only in anabolic processes but also their catabolism can contribute to energy metabolism^18^. For example, arginine catabolism directly yields ATP and is tightly regulated^19^, while other amino acids can contribute to pH or redox homeostasis, saving ATP-costly alternatives^20^. It is, however, not known whether proteome constraints guide choices in non-sugar substrates, e.g., amino acids.

We therefore developed pcLactis, a proteome-constrained genome-scale metabolic model of *L. lactis*. We first updated a published genome-scale metabolic model for *L. lactis* MG1363^21,22^ mostly by adding transport capacities and gene protein reaction associations (Supplementary Data 1). Second, we added the gene expression processes, including transcription, stable RNA cleavage, mRNA degradation, tRNA modification, rRNA modification, tRNA charging, ribosomal assembly, translation, protein maturation and assembly, and protein degradation (Fig 1a). By integrating the two reconstructions, we obtained pcLactis, which accounts for 725 protein-coding genes and 81 RNA genes in *L. lactis* MG1363. According to the PaxDb database^23^, pcLactis accounts for approximately 60% of the total proteome by mass (Supplementary Data 2). The model’s proteome therefore includes 40% unmodelled protein of average amino acid composition. We constrain metabolic fluxes by enzyme levels, following standard enzyme kinetics; Enzyme levels follow from mass balancing synthesis rate and degradation and dilution by growth (Fig 1b). Total protein synthesis rates are constrained by ribosomal translation capacity and by a total proteome constraint, i.e., a maximum size of the proteome. We use inactive enzyme, again with average amino acid composition, to fill up the total proteome in cases of low enzymatic activity, e.g. at low growth rates. Inactive enzyme can be seen as the excess capacity of the total modelled proteome.

**Fig 1.**
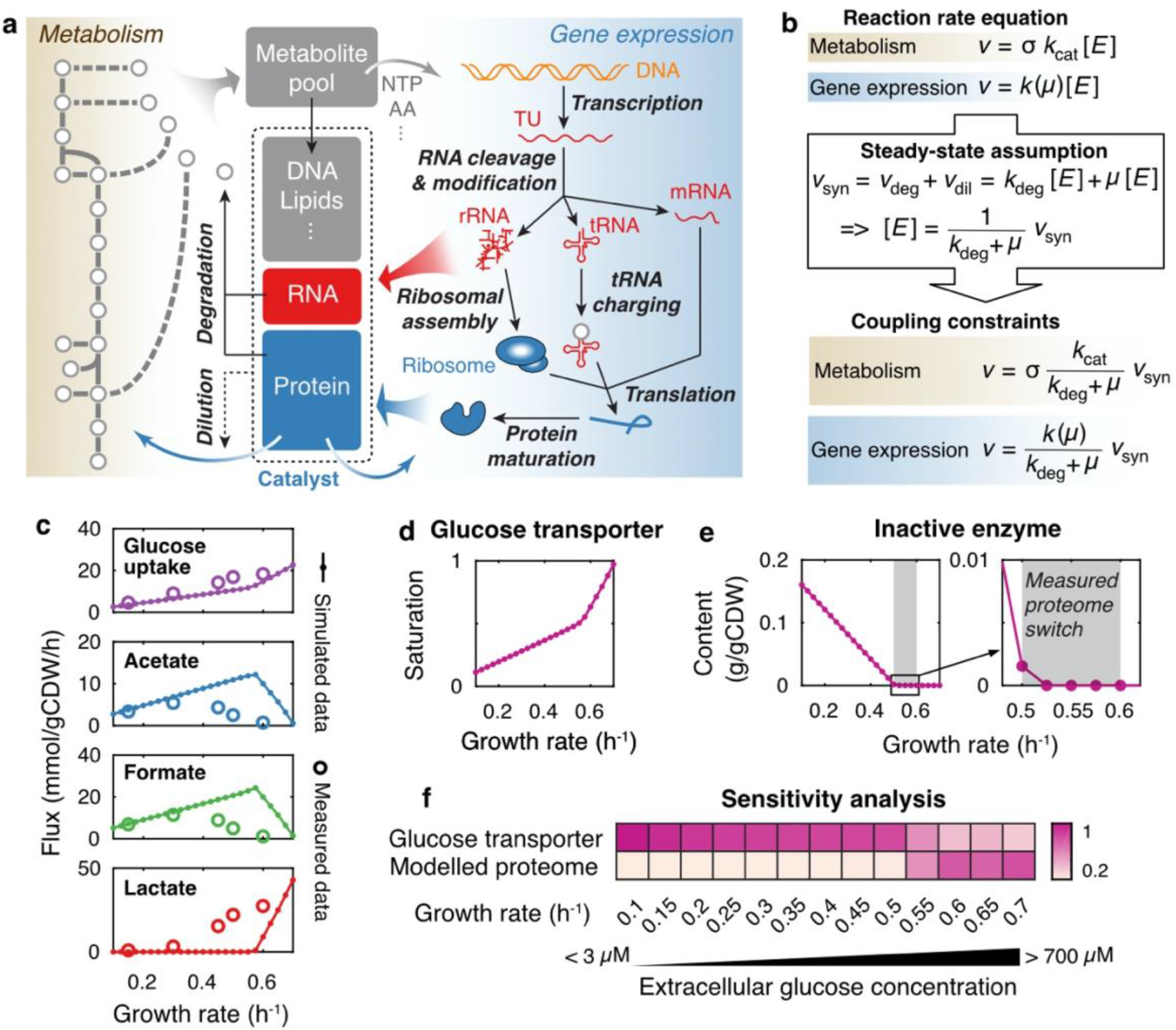
Overview of pcLactis and simulations of glucose-limited conditions. **a,** The model explicitly accounts for reactions of metabolism and gene expression processes. Metabolic reactions produce metabolites and energy for not only biomass formation but also gene expression processes. The reactions of gene expression, on the other hand, synthesize RNA and proteins, which catalyse reactions of metabolism and gene expression processes as machineries or enzymes. Besides, pcLactis accounts for degradation of mRNA and proteins as well as dilution of biomass constituents during cell division. **b,** Coupling constraints in pcLactis. The coupling constraint allows for relating the reaction rate to the synthesis rate of its catalyst based on the reaction rate equation and steady-state assumption, where turnover rates *k*_cat_ of metabolic enzymes, catalytic rates *k* of machineries and degradation constants *k*_deg_ of the catalysts are needed. **c,** Simulated exchange fluxes compared with experimentally measured data^3^. **d,** Simulated saturation of glucose transporter. **e,** Simulated inactive enzyme. The inactive enzyme is the sum of the enzymes that are synthesized but do not carry fluxes. The production of the inactive enzyme indicates that total proteome is not constrained. The grey area represents where the proteome switch occurs in experiments^3^, which is between 0.5 and 0.6 h^-1^. **f,** Sensitivity analysis for glucose transporter and modelled proteome at different growth rates. Colour represents the sensitivity score. A higher score indicates a greater impact of a given increase in the constraint on growth rate.

We used published proteomics and flux data from chemostats for model evaluation, and simulated glucose-limited conditions by minimizing the glucose concentration at a fixed specific growth rate with an upper bound on the expression of the glucose transporter. The model predicted the glucose uptake rates well (Fig 1c) and indicated-as expected-increased saturation of the glucose transporter with growth rate (Fig 1d). Additionally, pcLactis predicted overflow metabolism, i.e., a metabolic switch from mixed acid to lactic acid fermentation, corresponding to simultaneous changes at the proteome level (Fig S1a), but at a much higher growth rate than experimentally observed (Fig 1c). However, experimental evidence showed that the metabolic switch in *L. lactis* does not relate to considerable proteome changes^3^ and therefore the switch predicted by pcLactis might probably not reflect the true reason for overflow metabolism; The metabolic switch rather involves enzyme kinetics that are beyond the current stoichiometric model (see^24^ for a possible explanation). More importantly, the model predicted that at a dilution rate (D) higher than 0.5 h^-1^, the fraction of inactive enzyme becomes zero (Fig 1e). This means that at that point all proteins are maximally active and any flux change can only be brought about by changes in protein levels, not enzyme saturation. The proteomics data showed genome-wide protein reallocation when growth rate increased beyond that point^3^, validating the model prediction.

To identify the “active” constraints, i.e., constraints that limit the growth rate in pcLactis, we developed a sensitivity analysis for proteome-constrained models. Glucose transport expression was an active constraint at low growth rates and its sensitivity dropped at the moment the inactive protein reached zero (Fig 1f); At that point, the total proteome also became limiting. This reflects the increased demand in proteome resource at high growth rates, both for metabolic fluxes and protein translation machinery (Fig S1a). Thus, around D = 0.5 h^-1^, the model switched from glucose-limited to combined glucose and proteome limited.

In the model, the transition to proteome limited growth was reflected in amino acid metabolism, not overflow metabolism. However, including amino acid uptake in pcLactis was challenging as the amino acid consumption could not be constrained by the model due to insufficient data of expression and kinetics for amino acid transport systems. This would thus lead to an overestimation of uptake and growth rate when setting free exchange rates of all 20 amino acids to mimic the medium of *L. lactis*, where amino acids are mostly in excess. However, published amino acid data^3^ showed that apart from aspartate and glutamate all the detected amino acids were taken up linearly with growth rate (Fig 2a). We therefore imposed growth rate-dependent upper bounds on the uptake rates of all amino acids based on these measurements. To distinguish anabolism and catabolism, we compared the uptake of amino acids with their tRNA charging flux, which represents the flux towards protein synthesis. To analyse the results, we performed a so-called scaled reduced cost analysis, which is a sensitivity analysis of the uptake bounds on the growth rate (Fig 2b).

**Fig 2.**
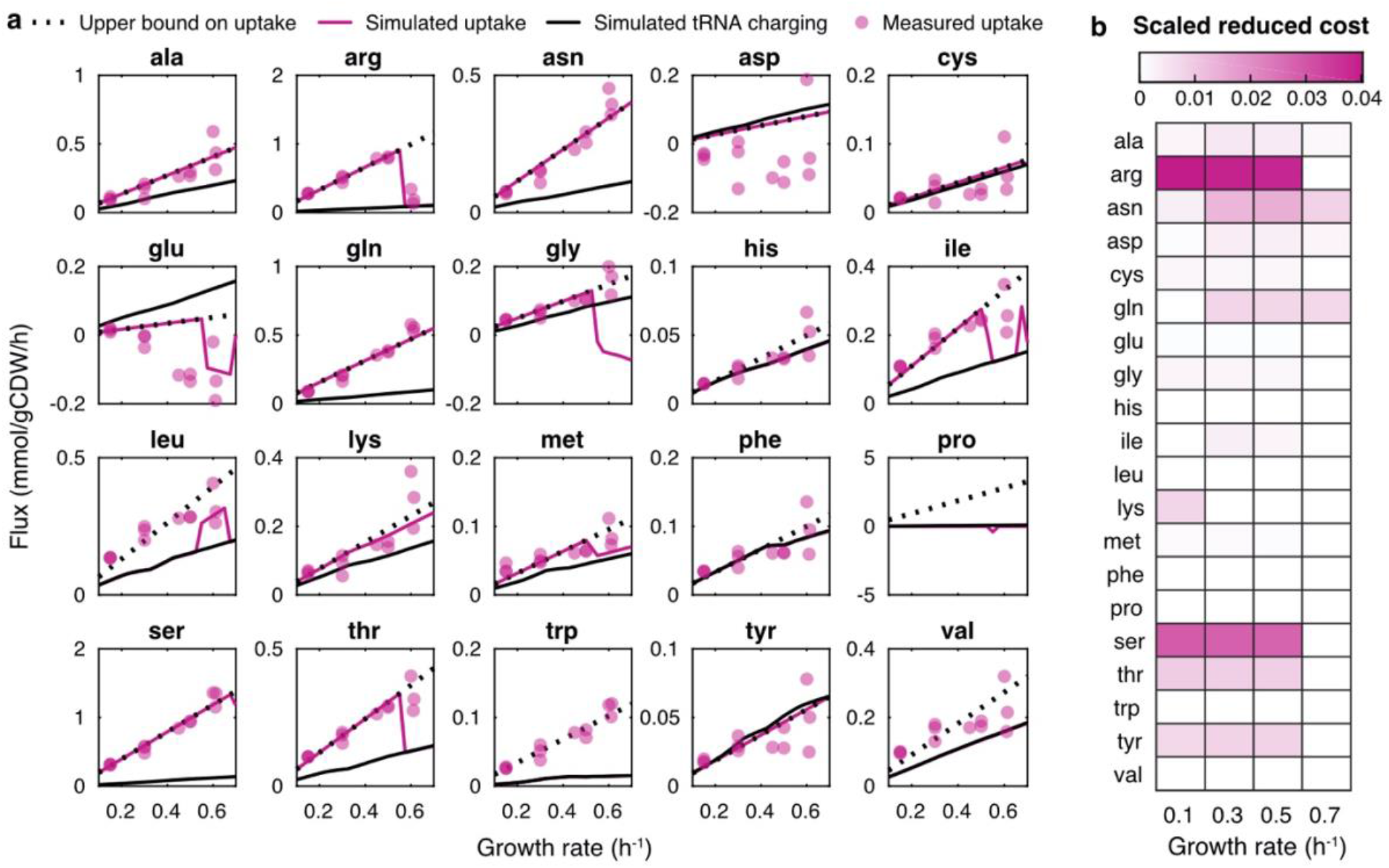
Amino acid analysis with pcLactis shows importance of amino acid catabolism for growth as a function of active constraints. **a,** Simulated fluxes of amino acids compared with experimentally measured data. For each amino acid, simulated fluxes of uptake and tRNA charging are displayed together with the measured uptake flux. The upper bound on the uptake is also displayed, which is the linear trendline of the experimentally measured data if the amino acid is consumed rather than being secreted. The measured data are from the study^3^. **b,** Scaled reduced cost analysis for uptake rates of amino acids at different growth rates. Colour represents the scaled reduced cost value. A higher value indicates a greater impact of a given increase in the amino acid uptake on growth rate.

The predicted uptake fluxes of most amino acids follow the upper bound set by the experimental data. Many of those amino acids were overconsumed and thus catabolized, in agreement with model predictions (Fig 2a). Based on the scaled reduced cost analysis, their catabolism contributed to growth rate (the mechanistic basis of many of those were previously analysed for *Lactobacillus plantarum*^25^). Those amino acids with little impact on growth rate, based on the scaled reduced cost analysis, are taken up by *L. lactis* according to protein synthesis demand. Thus, pcLactis can explain overconsumption of specific amino acids according to growth rate optimisation. This is interesting, as many of the catabolic products contribute to flavour formation in food fermentations. Leucine, tryptophan and to a lesser extent valine were exceptions (Fig 2a). In *L. plantarum* leucine and valine catabolism contributes to the redox balance, which requires a NADP-dependent glutamate dehydrogenase^25^; *L. lactis*, however, lacks this enzyme^26^. The reason for overconsumption of these amino acids thus remains to be unravelled.

When the total proteome becomes an active constraint at a growth rate around 0.5 h^-1^, we observe a drop in several amino acid overconsumptions in pcLactis. This is also reflected in a drop in reduced cost of the respective amino acids (Fig 2), indicating that the metabolic benefit was not high enough for the now constraining protein investment in the catabolic pathway. Experimentally, not all amino acids dropped at this point, although data is more variable at the highest growth rate. However, the most pronounced change in flux was observed for arginine, the amino acid that also had the highest (drop in) reduced cost. Arginine is also the only amino acid whose catabolism directly yields ATP. The predicted changes in the fluxes of catabolic products ornithine and ammonium (Fig S1b), and the concomitant changes in protein levels of the pathway (Fig S1a) together with experimental observations^3,19^, confirm that pcLactis correctly captured the switch in arginine catabolism.

We noted that the arginine switch preceded the onset of overflow metabolism (lactate) in pcLactis (Fig S1c). We constructed a small model to analyse this switching behaviour, expecting that protein efficiency, i.e. ATP produced per protein mass per time, was key^13^. We defined three independent ATP-producing pathways, glycolysis pathway with mixed acid fermentation or with lactate formation, and arginine catabolism (Fig 3a), and estimated ATP yield, protein cost and protein efficiency (Fig 3b) for each pathway using pcLactis. With these parameters, we formulated a linear program to maximize ATP production flux subject to total proteome constraints with flux bounds (Fig 3c). We varied glucose uptake fluxes and found three distinct phases: when the total proteome is not constrained both mixed acid fermentation and arginine catabolism are used (phase A, Fig 3d), as mixed acid fermentation has the highest ATP yield and arginine provides extra ATP at no protein burden. Once the total proteome is constrained (phase B, Fig 3d), the flux through arginine catabolism goes down due to its lowest protein efficiency. When arginine catabolism is completely inactive, mixed acid fermentation is traded in for lactate formation (phase C, Fig 3d), as its protein efficiency is lower than lactic acid fermentation. Furthermore, the small model reproduced the sensitivity results in terms of the constraints on glucose and arginine uptake (Fig 3d). Thus, the small model captures the behaviour of the full model, reinforcing earlier theoretical results that the number and nature of the active constraints determine behaviour, irrespective of the size of the network^24^.

**Fig 3.**
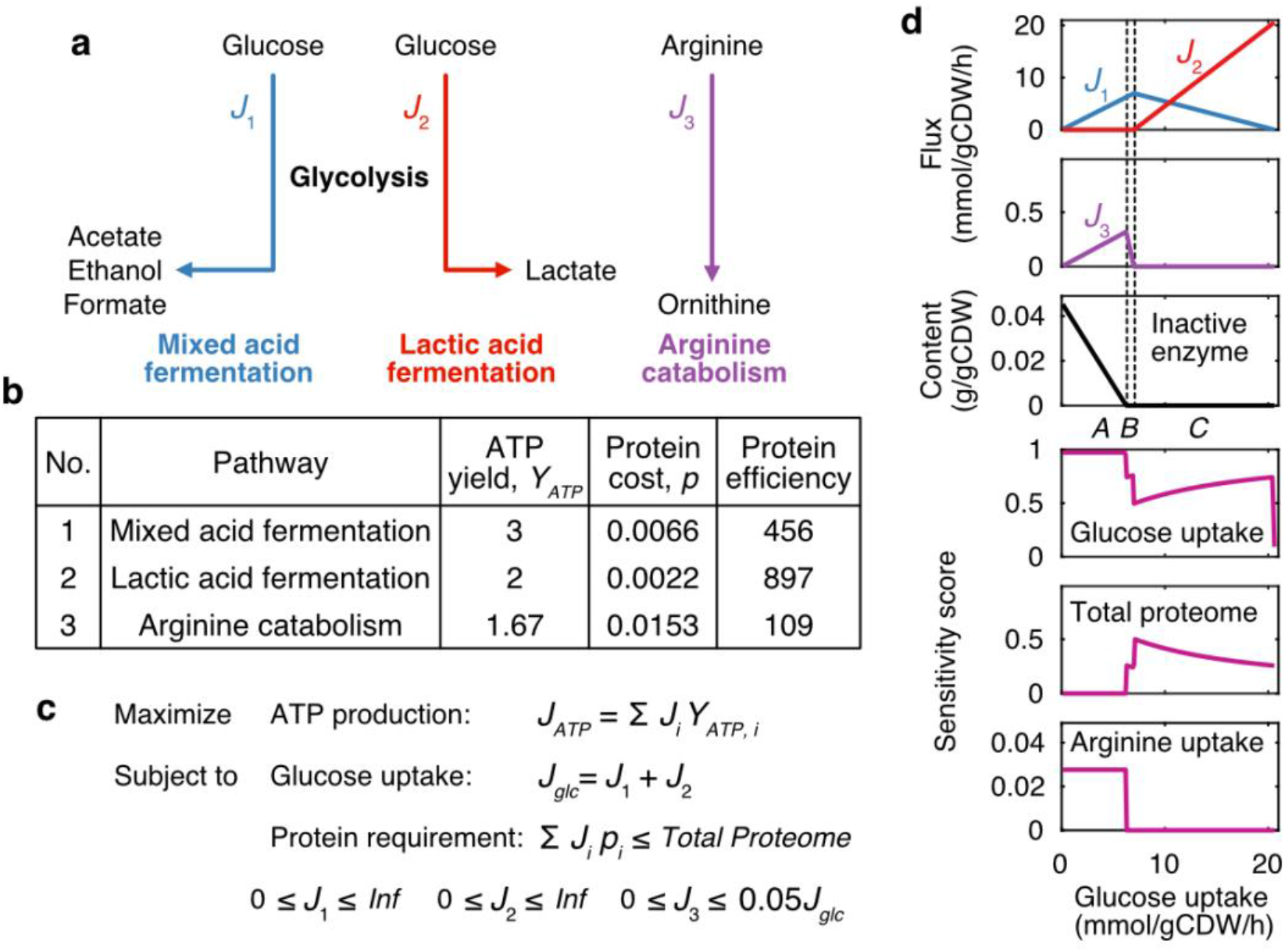
The switches between ATP yielding pathways in *L. lactis* can be explained from a protein efficiency point of view. **a,** A small model derived from pcLactis to investigate primary ATP-producing pathways in *L. lactis*. The small model consists of three independent pathways, including glycolysis pathway followed by mixed acid fermentation, glycolysis pathway followed by lactic acid fermentation and arginine catabolism. **b,** Parameters of ATP-producing pathways inferred from pcLactis. ATP yield is ATP generated per glucose (for pathway 1 and 2) or arginine (for pathway 3) consumed. Protein cost is protein mass (unit: g/gCDW) required per substrate flux (unit: mmol/gCDW/h) through the pathway. Protein efficiency is ATP generated per protein mass per time with the unit of mmol of ATP per gram of protein per hour. **c,** Linear programming to solve the small model. The objective function is to maximize the total ATP production flux *J_ATP_*, which is the sum of ATP production fluxes of three pathways calculated by substrate uptake flux *J_i_* times ATP yield *Y_ATP,i_*. The model is subject to two constraints. One is the total glucose uptake flux *J_glc_*, which is the sum of the glucose fluxes towards pathway 1 *J*_1_ and 2 *J*_2_. The other is the total proteome constraint, which means that the sum of protein requirements of all pathways should be not greater than the proteome allocation in the model. For each pathway, the protein requirement is calculated by the protein cost *p_i_* times substrate flux *J_i_*. Besides, there are lower and upper bounds on the uptake flux of each pathway. Specifically, *J*_1_ and *J*_2_ are unlimited while *J*_3_ has a limited upper bound, which is assumed to be linearly correlated with total glucose uptake flux *J_glc_* based on experimental observation. **d,** Small model simulations for a range of glucose uptake flux. The top three plots show that with increasing glucose uptake flux the decline in arginine uptake occurs firstly once the inactive enzyme disappears, and subsequently the switch from mixed to lactic acid fermentation. The inactive enzyme is the difference between the total protein requirements and the total proteome. The bottom three plots show sensitivities of glucose uptake, total proteome and arginine uptake to the ATP production.

Since pcLactis uses optimisation, we investigated if the predicted high sensitivities for glucose and arginine metabolism would provide targets for fitness improvements. For the simulations we used previously experimentally determined constraints on the uptake of amino acids^3^, which could be suboptimal under glucose-limited chemostat conditions. We therefore compared wild type *L. lactis* MG1363 with a mutant strain (designated 445C1) that evolved from MG1363 under glucose-limited chemostat conditions at D = 0.5 h^-1^, which harbours a point mutation in the global carbon catabolite repression regulator (CcpA)^27^. This CcpA mutant shows a twofold increase of mixed acid fermentation at the endpoint of the laboratory evolution experiment, while fermentation towards lactate is decreased^27^. This change in metabolism is consistent with the prediction that the total proteome constraint is not yet active at 0.5 h^-1^, as that would provide a driving force towards lactate formation (Fig 3).

To further validate predictions, we assessed protein allocation and amino acid metabolism of wild type and CcpA mutant, not studied before. We therefore re-cultivated both strains in glucose-limited chemostats at D = 0.5 h^-1^ and compared them at the proteome and metabolic level (Fig 4a). Principal component analysis of the proteomics data confirmed reproducibility (Fig S2). Apparent catabolic and total carbon balances were between 89 and 100% (Supplementary Data 3). We also found increased mixed acid and decreased lactic acid fermentation for the CcpA mutant compared to wild type (Fig S3), in agreement with the published data^27^.

**Fig 4.**
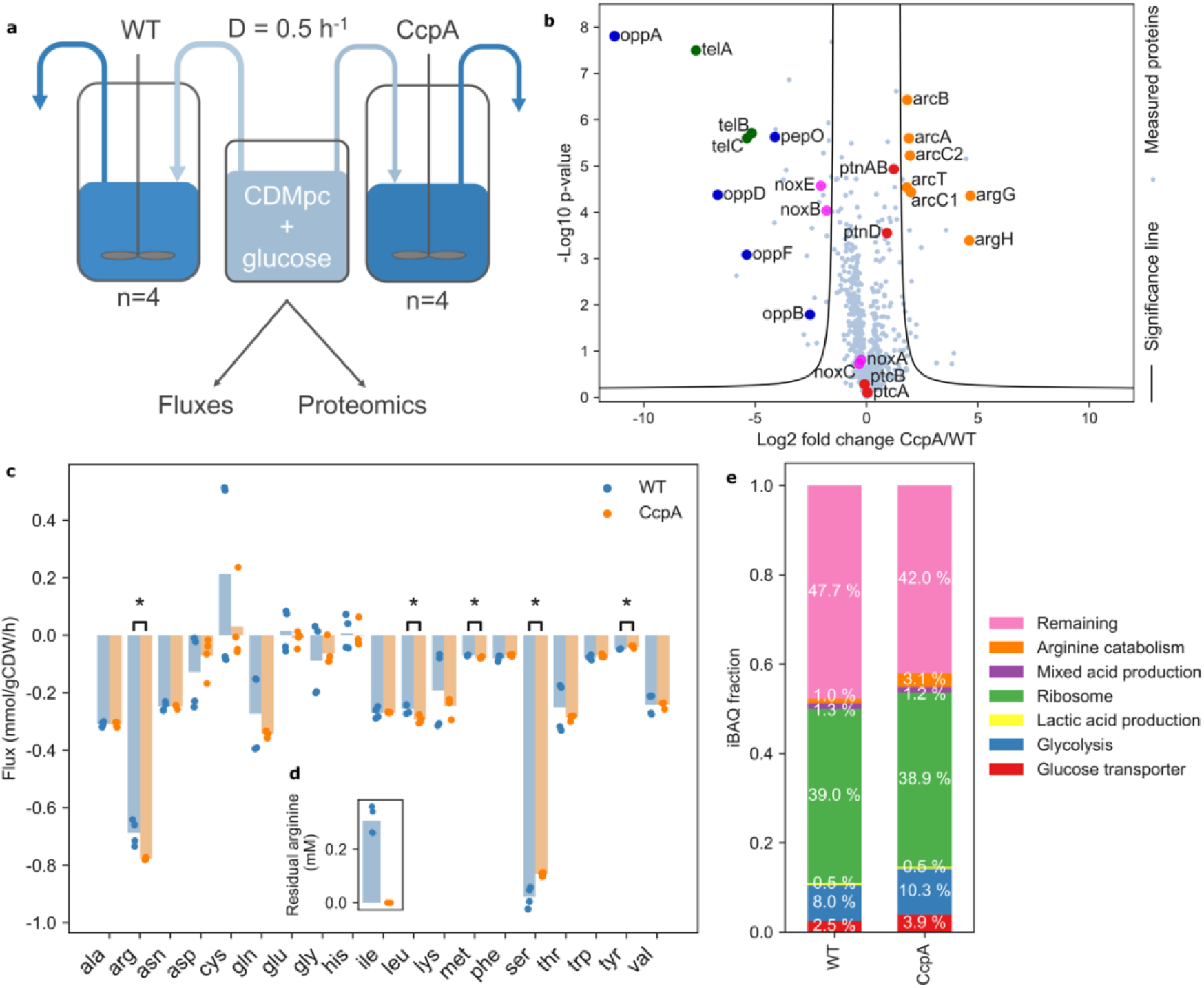
Proteome of the CcpA mutant, which has a higher fitness than wild type *L. lactis*, is changed in the direction of optimality. **a,** Overview of experiments for comparing wild type *L. lactis* MG1363 and CcpA mutant 445C1 in glucose-limited chemostats at D = 0.5 h^-1^. Samples were taken for dry weight measurements, external metabolite and amino acid analysis and proteome measurements. **b,** Protein fold changes of the CcpA mutant over wild type. Log2 fold changes were calculated from LFQ values *p* values were calculated based on 2-sided two sample t-tests with a FDR threshold of 0.05 and S_0_ equal to 1^40^. Proteins outside the significance lines (FDR = 0.05, S_0_ = 1) are significantly changed in expression. Example sets of proteins are highlighted and are labeled with their gene names. **c,** Amino acid fluxes (mmol/gCDW/h) and **d,** Residual concentration (mM) of arginine. The arginine uptake flux is increased to such extent that the residual arginine concentration became undetectable in the CcpA mutant. Data points represent the individual chemostats; bars are average values. **e,** Protein fractions. Protein fractions were obtained by normalizing iBAQ values by the total sum of all values. The obtained protein fractions were then averaged over the four biological replicates for visualization. The fractions glucose transport, glycolysis, arginine pathway and remaining fraction are significantly different between the two strains (*p* < 0.001, calculated with two-sided two sample t-tests).

The proteomics data showed that arginine catabolism, including its uptake system are significantly upregulated in the CcpA mutant compared to the wild type (Fig 4b; Fig S4). Consequently, we found an increased arginine uptake flux in the CcpA mutant (Fig 4c), to the extent that no residual arginine was detected anymore in its supernatant (Fig 4d). Other amino acids revealed no clear changes (Fig 4c, Fig S5), even though some had nonzero reduced costs as well, likely because metabolism of these amino acids is not under control of CcpA^28^. Taken together arginine could be the most effective amino acid on optimizing fitness under the chosen conditions as predicted by our model.

We further found a significantly increased protein fraction of glucose transporters in the CcpA mutant (*p* < 0.001) (Fig 4e), confirming that glucose uptake capacity was growth-limiting in the wild type. Even though the total proteome is not limiting at D = 0.5 h^-1^, we did find changes in cytosolic protein fractions, notably the glycolysis fraction (Fig 4e), while other proteins, not used by pcLactis under glucose limitation, were downregulated, e.g., peptide digestion and NADH oxidases (Fig 4b, Fig S4). These may be part of the global catabolite repression effect of the CcpA mutation without large fitness impact. Alternatively, upregulation of glycolysis may relieve inhibition on glucose transport, such as the negative impact of fructose 1,6-bisphosphate on the PTS system, via HPr^29^.

We did not find a considerable change in the total protein membrane fraction (Table S1), suggesting that the membrane protein occupancy is independent of the tested conditions and, given the need to transport many nutrients, possibly maximally occupied. If fully occupied, an increase of one protein would go at the cost of another protein. In the CcpA mutant we found an overrepresentation of significantly changed membrane proteins (*p* = 0.030 compared to random distribution of significantly changed proteins over membrane and cytosolic protein fraction, Table S1), amongst which downregulation of unused transport systems such as for peptides (Opp operon in Fig 4b). The idea that the membrane is fully occupied with proteins would also explain why amino acid uptake is not higher in wild type. It is therefore anticipated to impose a constraint on total membrane proteins in future simulations when transporter kinetics are available.

In conclusion, we show that integration of flux and proteomics data with a proteome-constrained genome-scale model allows for advanced sensitivity analysis of growth in complex nutrient environments. These sensitivities are predictive for evolutionary change, providing deeper understanding of the driving forces that shape regulatory strategies. We find changes in sensitivity that reflect which constraints become active under different growth rate regimes. Rather than the shift between mixed acid and lactic acid fermentation, we show that arginine metabolism, the third ATP yielding pathway in *L. lactis* and many other bacteria, is subject to proteome-constraint driven protein reallocation, mediated by CcpA. And after all, also *L. lactis* appears to abide to the microbial growth laws^30^ of cellular resource allocation.

## Methods

### Proteome-constrained model construction

The detailed construction procedure is described in the Supplementary Note. Firstly, we updated the existing genome-scale metabolic model of *L. lactis* MG1363^21,22^ in terms of transport reactions and gene-protein-reaction (GPR) associations by using MetaDraft (http://doi.org/10.5281/zenodo.2398336), a tool that reconstructs genome-scale metabolic networks based on previous manually curated ones by homology between genes, and subsequently manual curation. For non-spontaneous reactions with missing GPRs, we assigned a “dummy” protein as their catalysts to eliminate potential bias toward using them in simulations. Then we split reactions with isozymes into multiple reactions that each is catalysed by one isozyme, and reversible reactions into forward and reverse direction. In addition to updating and reformulating metabolic reactions, we formulated reactions for transcription, stable RNA cleavage, mRNA degradation, tRNA modification, rRNA modification, tRNA charging, ribosomal assembly, translation, protein maturation, protein assembly, enzyme formation, and protein degradation. Additionally, we formulated dilution reactions for RNA and enzymes to represent their dilution to daughter cells during cell division. Lastly, we modified the biomass equation of the metabolic model, i.e., removed protein and RNA from the equation as they were represented by dilution reactions, and added an unmodelled protein to account for all the other proteins that are not synthesized by the model. The genome-scale metabolic model was updated using CBMPy (https://doi.org/10.5281/zenodo.3485023), while pcLactis was constructed using the COBRA toolbox^31^.

### Constraints and kinetics parameters

In addition to classical constraints of GEMs, e.g., mass balance and bounds on reaction rates, two protein constraints are imposed including a fixed bound on the total modelled proteome and an upper bound on the abundance of glucose transporter (Supplementary Note). The fraction of modelled proteome was estimated according to the PaxDb database^23^, where abundance of each protein is collected. The data is however available only for *L. lactis* IL1403, and thus we performed BLASTp for mapping protein IDs between *L. lactis* IL1403 and MG1363. As a result, we obtained a list of proteins in *L. lactis* MG1363 with available abundance data (Supplementary Data 2). It should be noted that we filtered out the proteins of blocked reactions that would never carry fluxes in the model.

Besides, pcLactis accounts for coupling constraints that relate enzymes and machineries to their catalytic functions. In pcLactis such constraints are imposed for coupling enzymes to metabolic reactions, RNA polymerase to transcription reactions, ribosomes to translation reactions, and so on. All the coupling constraints are detailed in the Supplementary Note. In order to determine coefficients in the coupling constraints, we automatically retrieved turnover rates of metabolic enzymes from the BRENDA database^32^ using the GECKO toolbox^12^ but also manually adjusted some according to literatures. Besides, we estimated catalytic rates of gene expression machineries, including ribosome, RNA polymerase, mRNA and tRNA, in the same way as done for ME-Model of *E. coli*^9^ based on reported data^20,33^ for *L. lactis*. Detailed description can be found in the Supplementary Note.

### Simulations with pcLactis for anaerobic glucose-limited conditions

Since growth rate is integrated into coupling constraints in linear programming, it should be used as input for simulations. Therefore, we used a binary search workflow to obtain the minimal extracellular glucose concentration that leads to a feasible solution for any given growth rate to simulate glucose-limited conditions. This approach is based on the Michaelis-Menten equation with a fixed upper limit on the concentration of the glucose transporter:

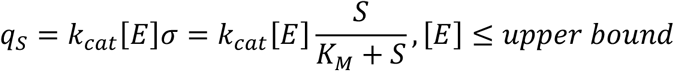

in which *q_S_* is the glucose uptake rate, *k_cat_* is the turnover rate of the glucose transporter adopted from *E. coli*^34^, [*E*] is the concentration of the glucose transporter, *σ* is the saturation of the glucose transporter, *S* is the extracellular glucose concentration and *K_M_* is the Michaelis constant obtained from the study^27^. The upper bound on the concentration of the glucose transporter was estimated using pcLactis, i.e., the minimal concentration that gives a feasible solution for the maximal growth rate. According to the Michaelis-Menten equation, searching for the minimal extracellular glucose concentration is equal to searching for the lowest saturation *σ*, which is equivalent to minimizing the glucose uptake rate when the upper bound of the concentration of the glucose transporter is hit.

The binary search solved successive individual linear programs, which maximizes the production of the dummy protein subject to stoichiometric and coupling constraints. Lower and upper bounds of some reactions were also constrained. Notably, amino acid exchange fluxes were constrained by growth rate-dependent upper bounds based on experimental data^3^. Besides, biomass dilution reaction, which represents the dilution of other biomass components than modelled protein and RNA, was fixed according to the growth rate. Due to the anaerobic condition we blocked two reactions, i.e., oxygen exchange reaction and pyruvate oxidase reaction. In addition, we blocked one alcohol dehydrogenase reaction catalysed by the isozyme llmg_0955, and two glucose transport reactions due to low protein levels^3^. Simulations were solved using Soplex 4.0.0 (https://soplex.zib.de/).

### Small model and simulations

The small model was extracted from pcLactis, which contains three pathways including glycolysis pathway with mixed acid fermentation, glycolysis pathway with lactic acid fermentation and arginine catabolism.

We estimated ATP yield, protein cost and protein efficiency for each pathway. Firstly, we identified the flux distribution of each pathway by solving a linear program with the metabolic part of pcLactis. For each simulation, the upper bound on ATP maintenance reaction was set free in order to account for ATP consumption. For mixed acid fermentation, the linear program is to maximize ATP maintenance reaction subject to a fixed glucose uptake rate of 1 mmol/gCDW/h. For lactate acid fermentation, the linear program is to maximize lactate production subject to a fixed glucose uptake rate of 1 mmol/gCDW/h and ATP maintenance reaction rate of 2 mmol/gCDW/h. For arginine catabolism, the linear program is to maximize ATP maintenance reaction subject to a fixed arginine uptake rate of 1 mmol/gCDW/h. Accordingly, we calculated ATP yield for each pathway, i.e., the flux of ATP maintenance reaction over uptake rate of substrate. Secondly, we estimated the protein cost for each pathway based on the flux distribution, which is the protein cost of each reaction in the pathway times the corresponding flux value. The protein cost of a reaction is molecular weight of the corresponding enzyme over its turnover rate^13^, and therefore can be extracted from pcLactis. Lastly, the protein efficiency of each pathway is ATP yield over protein cost.

With the parameters, we generated a linear program of fluxes through the three pathways, which is to maximize ATP production flux subject to constraints on uptake fluxes of substrates, i.e., glucose and arginine, and total proteome. We performed the simulations for a wide range of glucose uptake fluxes.

### Scaled reduced cost analysis

We performed the scaled reduced cost analysis^25^ for uptake rate of each amino acid on growth rate with pcLactis. The scaled reduced cost *R_i_* of the growth rate *μ_i_* with respect to the uptake rate of an amino acid *q_i_* is calculated as: *R_i_* = (Δ*μ*/Δ*q*)(*q_i_*/*μ_i_*). We imposed a small increase in the uptake rate of each amino acid Δ*q* = 0.01 to investigate the change in growth rate Δ*μ*.

### Sensitivity analysis

We performed sensitivity analysis for two constraints with pcLactis, i.e., the glucose transporter and modelled proteome, on growth rate. The sensitivity score *S_i_* of the growth rate *μ_i_* with respect to a given constraint *c_i_* is calculated as: *S_i_* = (Δ*μ*/Δ*c*)(*c_i_*/*μ_i_*). With pcLactis, we imposed a small increase Δ*c* = 0.01 in the glucose transporter and modelled proteome to investigate the change in growth rate Δ*μ* at different conditions. We also performed sensitivity analysis with the small model for glucose uptake, proteome allocated to the small model and arginine uptake using the small increase Δ*c* = 0.01 to investigate the change in ATP production rate *q_ATP_*, i.e., the sensitivity score *S_i_* is calculated as: *S_i_* = (Δ*q_ATP_*/Δ*c*)(*c_i_*/*q_ATP_*).

### Strains and media

*Lactococcus lactis* ssp. *cremoris* MG1363 is a plasmid cured derivative of strain NCDO712^35,36^. Strain 445C1 is a derivative of MG1363 isolated after prolonged cultivation in a chemostat^27^. Cultivation of strains were performed in chemically defined medium for prolonged cultivation (CDMpc)^27^, supplemented with 25 mM glucose as limiting carbon source at 30°C.

### Chemostat cultivation

Proteome, metabolite and amino acid samples were obtained from four steady state chemostats. Chemostat cultivation took place in 300 mL bioreactors with 270 mL working volume, under continuous stirring (using magnetic stirrers), while standing in 30°C water baths. The headspace was continuously flushed with 5% CO_2_ and 95% N_2_. pH was controlled at 6.5 using 2.5 M NaOH. The pH probe was calibrated before and after the preculture in the reactor. CDMpc with 25 mM glucose was added at a rate of 2.25 mL/min. Superfluous liquid was continuously removed from the top of the reactor to maintain a volume of 270 mL. This results in a dilution rate of 0.5 h^-1^. The flow rate of each medium pump was calibrated right before inoculating the reactor and checked again at the end of the experiment. The exact volume of liquid in each reactor was determined at the end of the experiment by weighing the reactor liquid.

To start up a chemostat culture, 5 mL CDMpc with 25 mM glucose was inoculated with glycerol stocks of the respective strains, taken from a −80°C. This was directly added to the reactor containing 265 mL fresh 25 mM glucose CDMpc. The reactor was operated in batch mode (no medium addition, pH regulated, effluent pump on, 30 °C, continuous stirring, headspace sparged) for 24 hours, allowing the cells to reach the stationary phase. This was verified by observing that the pH remained constant without addition of 2 M NaOH. After 24 hours the medium pump was switched on. The chemostat was operated for 8 volume changes before samples were taken. Samples were taken from the effluent of the reactor. Retention time in the effluent tube was <2 minutes. Sample tubes were kept on ice while sample was being collected. For each strain four replicate chemostats were cultivated.

### Sampling procedures and measurements

For extracellular metabolite concentration measurements (for both reactor broth and sterile medium), 2 mL samples were centrifuged at 27237 G for 3 minutes at 4 °C, the supernatant was filtered through a 0.22 μm polyether sulfone filter (VWR international) and stored at −20 °C until further analysis. Extracellular concentrations of lactate, acetate, formate, ethanol and glucose were determined by high-performance liquid chromatography (HPLC) as described previously^37^.

Extracellular concentrations of amino acids were determined by HPLC on a Shimadzu system with: LC-20AD pumps, DGU-14A degasser, SIL-10ADvp autosampler, CTO-10ASvp oven, SCL-10Avp system controller, and RF-10AXL fluorescence detector. Separation occurred on an Agilent Zebra Eclipse plus solvent saver 3.0×150×3.5 column over a period of 38 minutes per sample, with an isocratic flow of two eluents both at a flow rate of 0.64 mL/min. Eluent 1 had composition: 0.142 % w/v NaHPO_4_, 0.381% w/v Na_2_B_4_O_7_·10H_2_O, 0.0325% w/v sodium azide in deionized water (DI water); and eluent 2 had composition: 45% v/v methanol, 45% v/v acetonitrile in DI water. Samples were prepared by mixing 25 μL sample with 875 μL DI water, 25 μL Borate buffer (0.6% w/v boric acid, 0.4% w/v NaOH, pH 10.2), and 25 μL 1mM Norvaline as internal standard. For amino acid derivation 3 μL phthaldialdehyde (Sigma-Aldrich), i.e. OPA, was automatically added 3 minutes before sample injection (5 μL) into the column. Concentrations were determined by comparison of sample peak areas to those of a calibration curve identically run with representative amino acid concentrations.

Proteome samples were collected in low protein binding tubes, centrifuged at 27237 G for 3 minutes at 4°C. The supernatant was discarded, and the pellet was frozen in liquid N_2_ prior to storage at −20°C until further analysis.

For dry weight measurements, 10 mL broth was collected and filtered over a pre-dried (3 days, 60 °C) pre-weighed 0.2 μm cellulose nitrate filter (Whatman GmbH). The filter was then washed once with 10 mL demineralized water and dried (3 days, 60°C), before weighing again.

Fluxes (*q_i_* in mmol/gCDW/h) were calculated as: *q_i_* = *D* · (*C_i,supernatant_* – *C_i,medium_*)/*X_biomass_*, where D is the dilution rate (h^-1^), C the concentration of compound i (mmol/L) and X_biomass_ the biomass concentration (g/L).

### Proteome measurements

Proteins were isolated using the FASP method as described previously^38^. In short, cell pellets were dissolved in 100 mM TRIS pH 8 to a concentration of 7.5E8 cells/100 μl and lysed using a needle sonicator (MPE). Proteins obtained from 4.5E8 lysed cells were reduced with 15 mM dithiothreitol for 30 minutes at 45 °C and subsequently alkylated in 20 mM acrylamide for 10 minutes at room temperature under denaturing conditions (100 mM TRIS pH8 + 8 M Urea). The alkylated sample was transferred to an ethanol-washed Pall 3K omega filter (Sigma-Aldrich, The Netherlands) and centrifuged for 36 minutes at 12000 rpm. The filters were washed with 130 μl 50 mM ammonium bicarbonate and centrifuged again before overnight digestion with 100 μl 5 ng/μl trypsin (Roche) at room temperature. The digested peptides were eluted by centrifugation for 30 min, subsequent addition of 100 μl 1 ml/l HCOOH in water and again centrifugation for 30 min. pH was adjusted to pH 3 using trifluoroacetic acid.

MS/MS analysis was done as described previously^39^. Raw datafiles were analyzed using MaxQuant (version 1.6.1.0) and searched against the *L. lactis* MG1363 database (UniProt) and frequently observed contaminants. In addition to the standard settings, Trypsin/P with a maximum of two missed cleavages was set as the digestion mode, acrylamide modifications on the cysteines was set as a fixed modification, and methionine oxidation, protein N-terminal acetylation and asparagine or glutamine deamidation were set as variable modifications. A false discovery rate of 1% at protein level was allowed and the minimum required protein length was set at 7. At least two peptides were required for protein identification of which at least one peptide was required to be unique in the database. Identified proteins were quantified.

### Proteomics data analysis

Statistical analysis on the MaxQuant output was performed with Perseus version 1.6.2.1. Proteins were accepted when they were represented in at least three of the four biological replicates in at least one strain. For statistical analysis log10 transformed LFQ values were used and zero values were replaced by taking random values from a normal distribution with mean (measured values per biological sample −1.8) and variation (0.3 * standard deviation of the measured values per biological replicate) to make calculations possible. We then performed a 2-sided two sample t-tests using the log10 normalized LFQ intensity columns of CcpA mutant and wild type with a FDR threshold of 0.05 and S_0_ equal to 1^40^. Fold changes of CcpA mutant over wild type were calculated by dividing LFQ intensity columns of CcpA mutant by wild type. A principal component analysis of the LFQ data was used to assess the reproducibility of the data. Further analysis steps were done with Python. For fractions of protein groups, we used sums of intensity-based absolute quantitation (iBAQ) protein fractions, which were obtained by normalizing iBAQ values by the total sum of all values per biological replicate. Two-sided two sample t-tests were performed on biological groups of proteins between the wild type and CcpA mutant strain. Furthermore, a two-sided Fisher exact test was performed on the number of significantly changed proteins and unchanged proteins in the membrane and rest of the cell. For this, the protein fractions were first averaged over the biological replicates.

## Supporting information

Supplementary Note - pcLactis Construction

## Data availability

The mass spectrometry proteomics data have been deposited to the ProteomeXchange Consortium via the PRIDE^41^ partner repository with the dataset identifier PXD021956.

## Code availability

The model and codes are available online at https://github.com/SysBioChalmers/pcLactis.

## Acknowledgements

Y.C. and J.N. acknowledge funding from the European Union’s Horizon 2020 research and innovation program under Grant Agreement No. 686070. Y.C. and J.N. also acknowledge funding from the Novo Nordisk Foundation (grant no. NNF10CC1016517). E.v.P.-K., B.v.O., S.B., H.B., D.M. and B.T. acknowledge funding from the Netherlands Organisation for Scientific Research (grant no. ALWTF.2015.4). S.D., H.B., D.M. and B.T. acknowledge funding from the Top-sector Agri&Food (grant no. AF-15503).

## Author contributions

B.T., H.B. and J.N. conceived the study. Y.C. and E.v.P.-K. performed the modelling, simulations and data-analysis. B.v.O. and S.D. performed experiments. D.M. and S.B. contributed to the data-analysis. Y.C., E.v.P.-K. and B.T. wrote the manuscript. All authors contributed to manuscript editing.

## Reference

1. Goel, A., Wortel, M. T., Molenaar, D. & Teusink, B. Metabolic shifts: a fitness perspective for microbial cell factories. Biotechnol. Lett. 34, 2147–2160 (2012).

2. Basan, M. Resource allocation and metabolism: the search for governing principles. Curr. Opin. Microbiol. 45, 77–83 (2018).

3. Goel, A. et al. Protein costs do not explain evolution of metabolic strategies and regulation of ribosomal content: does protein investment explain an anaerobic bacterial Crabtree effect? Mol. Microbiol. 97, 77–92 (2015).

4. Chubukov, V., Gerosa, L., Kochanowski, K. & Sauer, U. Coordination of microbial metabolism. Nat. Rev. Microbiol. 12, 327–340 (2014).

5. Beg, Q. K. et al. Intracellular crowding defines the mode and sequence of substrate uptake by Escherichia coli and constrains its metabolic activity. Proc. Natl. Acad. Sci. 104, 12663–12668 (2007).

6. Molenaar, D., van Berlo, R., de Ridder, D. & Teusink, B. Shifts in growth strategies reflect tradeoffs in cellular economics. Mol. Syst. Biol. 5, 323 (2009).

7. de Groot, D. H. et al. The common message of constraint-based optimization approaches: overflow metabolism is caused by two growth-limiting constraints. Cellular and Molecular Life Sciences 77, 441–453 (2020).

8. Zhuang, K., Vemuri, G. N. & Mahadevan, R. Economics of membrane occupancy and respiro-fermentation. Mol. Syst. Biol. 7, 500 (2011).

9. O’Brien, E. J., Lerman, J. A., Chang, R. L., Hyduke, D. R. & Palsson, B. Genome-scale models of metabolism and gene expression extend and refine growth phenotype prediction. Mol. Syst. Biol. 9, 693 (2013).

10. Basan, M. et al. Overflow metabolism in Escherichia coli results from efficient proteome allocation. Nature 528, 99–104 (2015).

11. Nilsson, A. & Nielsen, J. Metabolic Trade-offs in Yeast are Caused by F1F0-ATP synthase. Sci. Rep. 6, 22264 (2016).

12. Sánchez, B. J. et al. Improving the phenotype predictions of a yeast genome-scale metabolic model by incorporating enzymatic constraints. Mol. Syst. Biol. 13, 935 (2017).

13. Chen, Y. & Nielsen, J. Energy metabolism controls phenotypes by protein efficiency and allocation. Proc. Natl. Acad. Sci. U. S. A. 116, 17592–17597 (2019).

14. Papadimitriou, K. et al. Stress Physiology of Lactic Acid Bacteria. Microbiol. Mol. Biol. Rev. 80, 837–90 (2016).

15. Kok, J. et al. The Evolution of gene regulation research in Lactococcus lactis. FEMS Microbiol. Rev. 41, S220–S243 (2017).

16. Teusink, B. & Molenaar, D. Systems biology of lactic acid bacteria: For food and thought. Curr. Opin. Syst. Biol. 6, 7–13 (2017).

17. Jensen, P. R. & Hammer, K. Minimal Requirements for Exponential Growth of Lactococcus lactis. Appl. Environ. Microbiol. 59, (1993).

18. Fernández, M. & Zúñiga, M. Amino Acid Catabolic Pathways of Lactic Acid Bacteria. Crit. Rev. Microbiol. 32, 155–183 (2006).

19. Crow, V. L. & Thomas, T. D. Arginine metabolism in lactic streptococci. J. Bacteriol. 150, 1024–1032 (1982).

20. Novák, L. & Loubiere, P. The metabolic network of Lactococcus lactis: Distribution of 14C-labeled substrates between catabolic and anabolic pathways. J. Bacteriol. 182, 1136–1143 (2000).

21. Flahaut, N. A. L. et al. Genome-scale metabolic model for Lactococcus lactis MG1363 and its application to the analysis of flavor formation. Appl. Microbiol. Biotechnol. 97, 8729–8739 (2013).

22. Verouden, M. P. H. et al. Multi-way analysis of flux distributions across multiple conditions. J. Chemom. 23, 406–420 (2009).

23. Wang, M., Herrmann, C. J., Simonovic, M., Szklarczyk, D. & von Mering, C. Version 4.0 of PaxDb: Protein abundance data, integrated across model organisms, tissues, and cell-lines. Proteomics 15, 3163–3168 (2015).

24. de Groot, D. H., van Boxtel, C., Planqué, R., Bruggeman, F. J. & Teusink, B. The number of active metabolic pathways is bounded by the number of cellular constraints at maximal metabolic rates. PLOS Comput. Biol. 15, e1006858 (2019).

25. Teusink, B. et al. Analysis of growth of Lactobacillus plantarum WCFS1 on a complex medium using a genome-scale metabolic model. J. Biol. Chem. 281, 40041–8 (2006).

26. Wels, M., Siezen, R., Van Hijum, S., Kelly, W. J. & Bachmann, H. Comparative genome analysis of Lactococcus lactis indicates niche adaptation and resolves genotype/phenotype disparity. Front. Microbiol. 10, 4 (2019).

27. Price, C. E. et al. Adaption to glucose limitation is modulated by the pleotropic regulator CcpA, independent of selection pressure strength. BMC Evol. Biol. 19, 15 (2019).

28. Zomer, A. L., Buist, G., Larsen, R., Kok, J. & Kuipers, O. P. Time-resolved determination of the CcpA regulon of Lactococcus lactis subsp. cremons MG1363. in Journal of Bacteriology 189, 1366–1381 (American Society for Microbiology Journals, 2007).

29. Deutscher, J., Francke, C. & Postma, P. W. How Phosphotransferase System-Related Protein Phosphorylation Regulates Carbohydrate Metabolism in Bacteria. Microbiol. Mol. Biol. Rev. 70, 939–1031 (2006).

30. Scott, M., Gunderson, C. W., Mateescu, E. M., Zhang, Z. & Hwa, T. Interdependence of cell growth and gene expression: origins and consequences. Science 330, 1099–102 (2010).

31. Heirendt, L. et al. Creation and analysis of biochemical constraint-based models using the COBRA Toolbox v.3.0. Nat. Protoc. 1 (2019). doi:10.1038/s41596-018-0098-2

32. Jeske, L., Placzek, S., Schomburg, I., Chang, A. & Schomburg, D. BRENDA in 2019: a European ELIXIR core data resource. Nucleic Acids Res. 47, D542–D549 (2019).

33. Beresford, T. & Condon, S. Physiological and genetic regulation of rRNA synthesis in Lactococcus. J. Gen. Microbiol. 139, 2009–2017 (1993).

34. Szenk, M., Dill, K. A. & de Graff, A. M. R. Why Do Fast-Growing Bacteria Enter Overflow Metabolism? Testing the Membrane Real Estate Hypothesis. Cell Syst. 5, 95–104 (2017).

35. Gasson, M. J. Plasmid complements of Streptococcus lactis NCDO 712 and other lactic streptococci after protoplast-induced curing. J. Bacteriol. 154, (1983).

36. Wegmann, U. et al. Complete genome sequence of the prototype lactic acid bacterium Lactococcus lactis subsp. cremoris MG1363. J. Bacteriol. 189, 3256–70 (2007).

37. Goel, A., Santos, F., Vos, W. M. de, Teusink, B. & Molenaar, D. Standardized assay medium to measure Lactococcus lactis enzyme activities while mimicking intracellular conditions. Appl. Environ. Microbiol. 78, 134–43 (2012).

38. Wiśniewski, J. R., Zougman, A., Nagaraj, N. & Mann, M. Universal sample preparation method for proteome analysis. Nat. Methods 6, 359–362 (2009).

39. Sotoca, A. M. et al. Quantitative Proteomics and Transcriptomics Addressing the Estrogen Receptor Subtype-mediated Effects in T47D Breast Cancer Cells Exposed to the Phytoestrogen Genistein. Mol. Cell. Proteomics 10, M110.002170 (2011).

40. Tusher, V. G., Tibshirani, R. & Chu, G. Significance analysis of microarrays applied to the ionizing radiation response. Proc. Natl. Acad. Sci. U. S. A. 98, 5116–5121 (2001).

41. Vizcaíno, J. A. et al. 2016 update of the PRIDE database and its related tools. Nucleic Acids Res. 44, D447–D456 (2016).

