## Supplementary Information for "Proteome constraints reveal targets for improving microbial fitness in nutrient-rich environments"

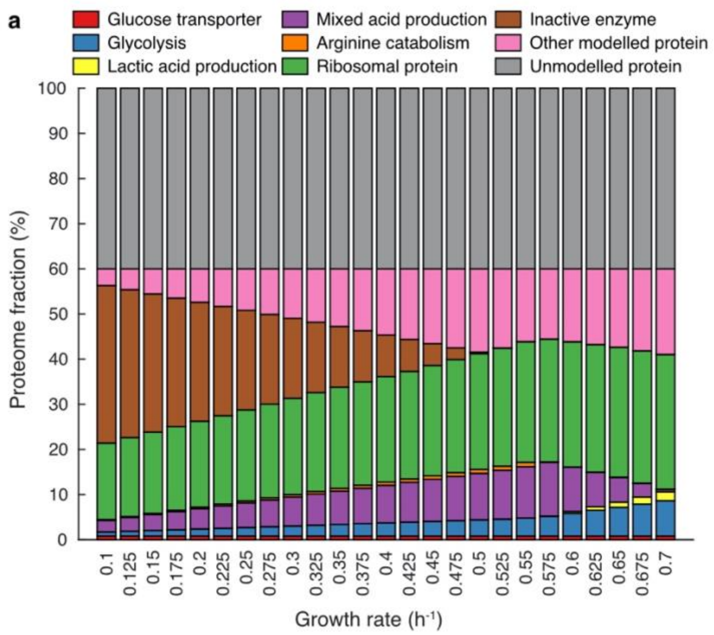


**Supplementary Fig. 1. Supplementary plots of simulations of glucose-limited conditions. a**, Simulated proteome allocation. **b,** Simulated exchanged fluxes of products related to arginine catabolism. **c,** Simulated arginine uptake fluxes versus lactate production fluxes in the range between growth rate of 0.53 and 0.58 $h^{-1}$.

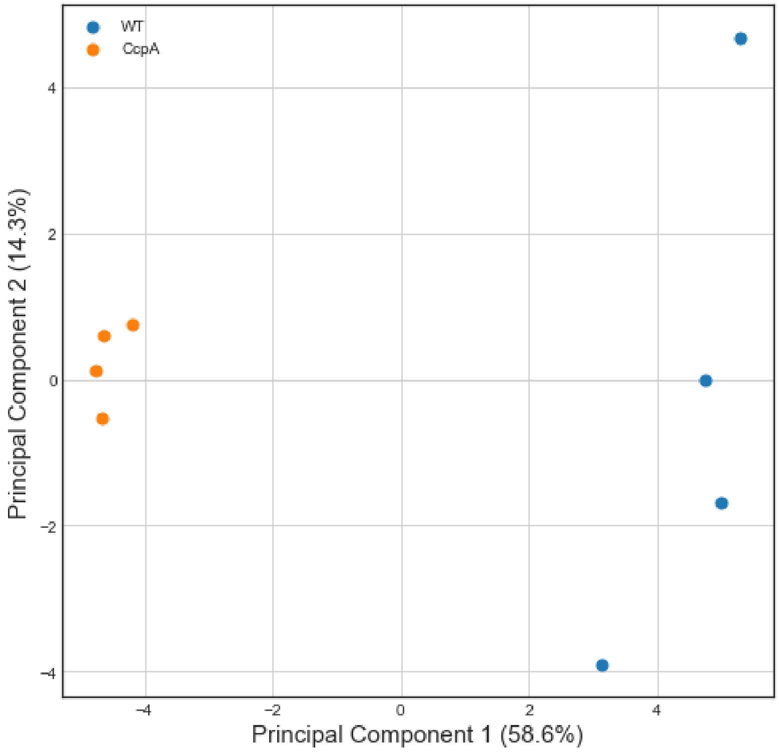


**Supplementary Fig. 2. Principal Component Analysis of Label Free Quantification protein data of the CcpA mutant 445C1 and wild type (WT) *L. lactis* MG1363.** The LFQ data was first log10 transformed. The four samples per strain cluster nicely together in individual groups.

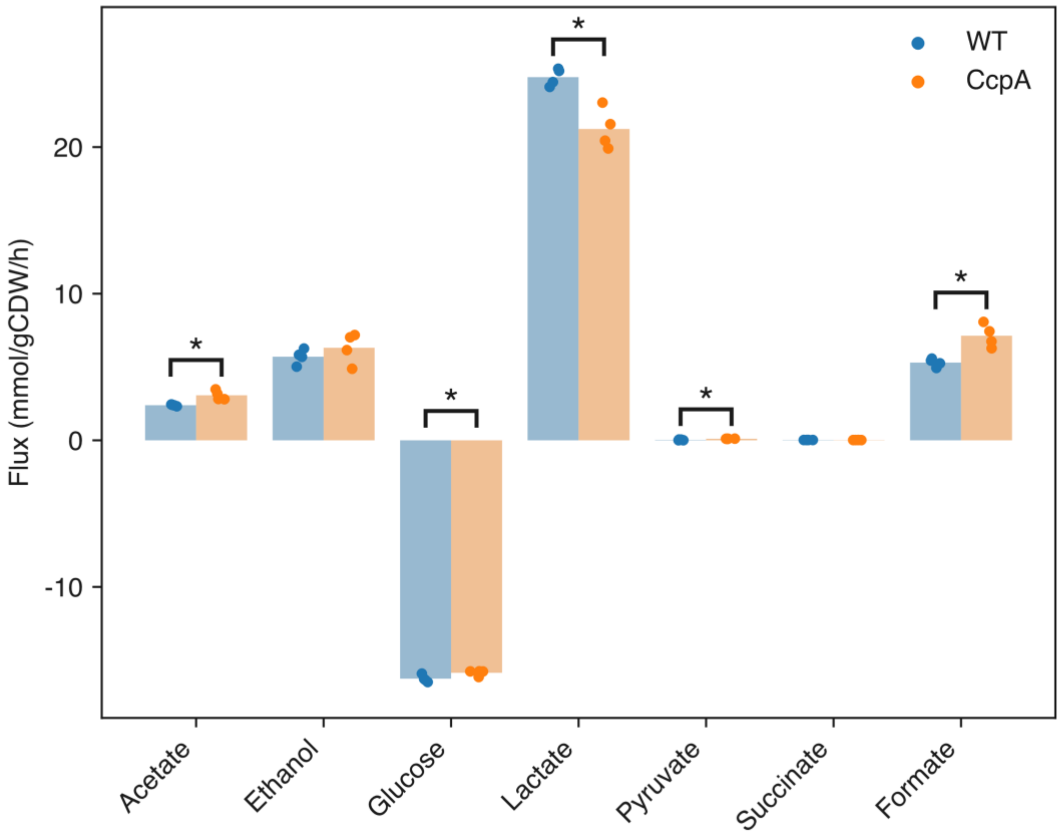


**Supplementary Fig. 3. Exchange fluxes for WT *L. lactis* MG1363 and CcpA mutant 455C1.** Data points represent the individual chemostats; bars are average values; * indicates a significant difference between WT and CcpA mutant, $p < 0.05$ for independent t-test.

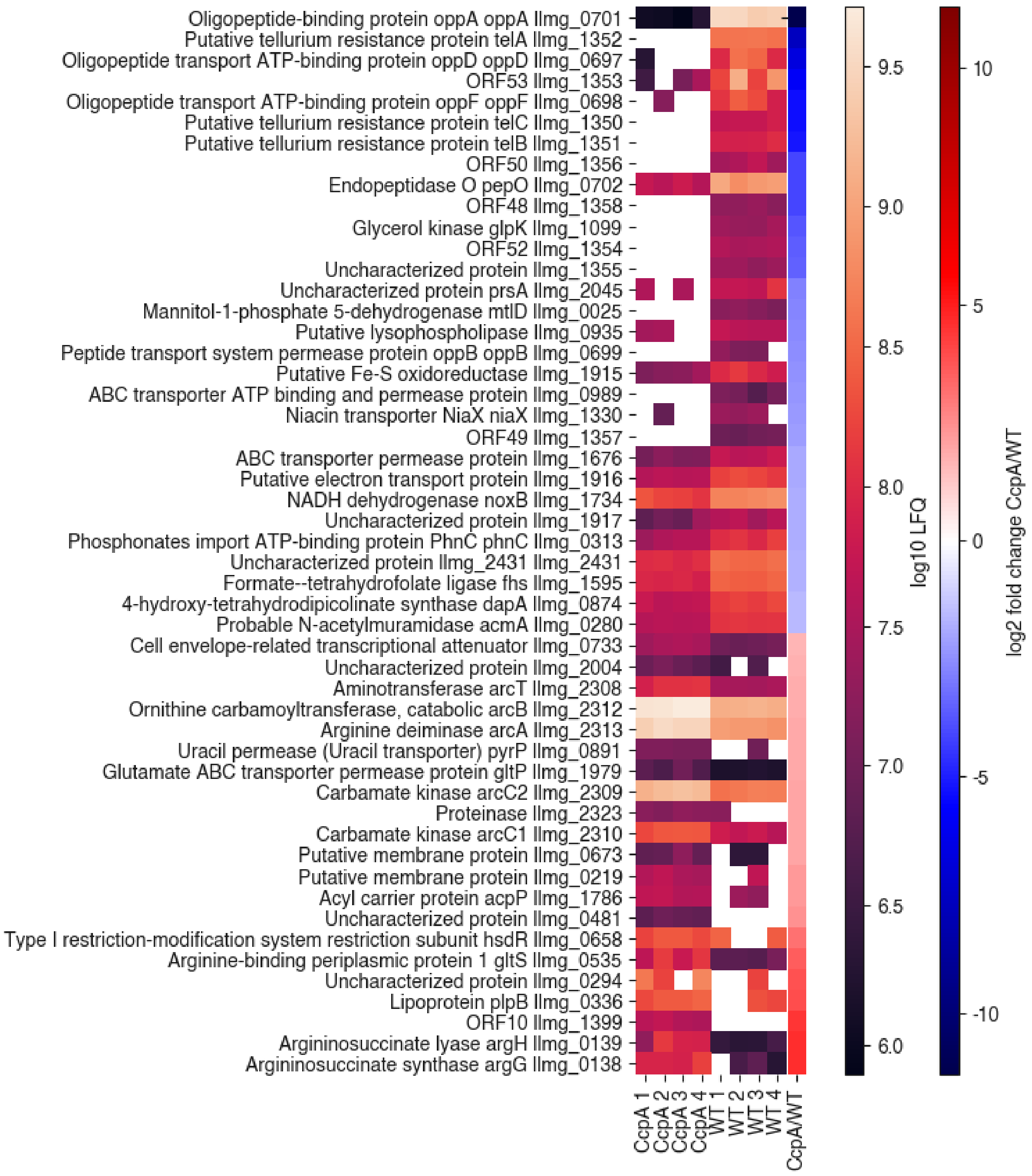


**Supplementary Fig. 4. Proteins that are significantly changed between WT *L. lactis* MG1363 and the CcpA mutant 445C1.** Pictured are log10 transformed LFQ data of CcpA mutant and WT *L. lactis* and log2 transformed fold changes CcpA mutant/WT. Proteins were ordered according to their log2 fold change value. NA values are indicated with white.

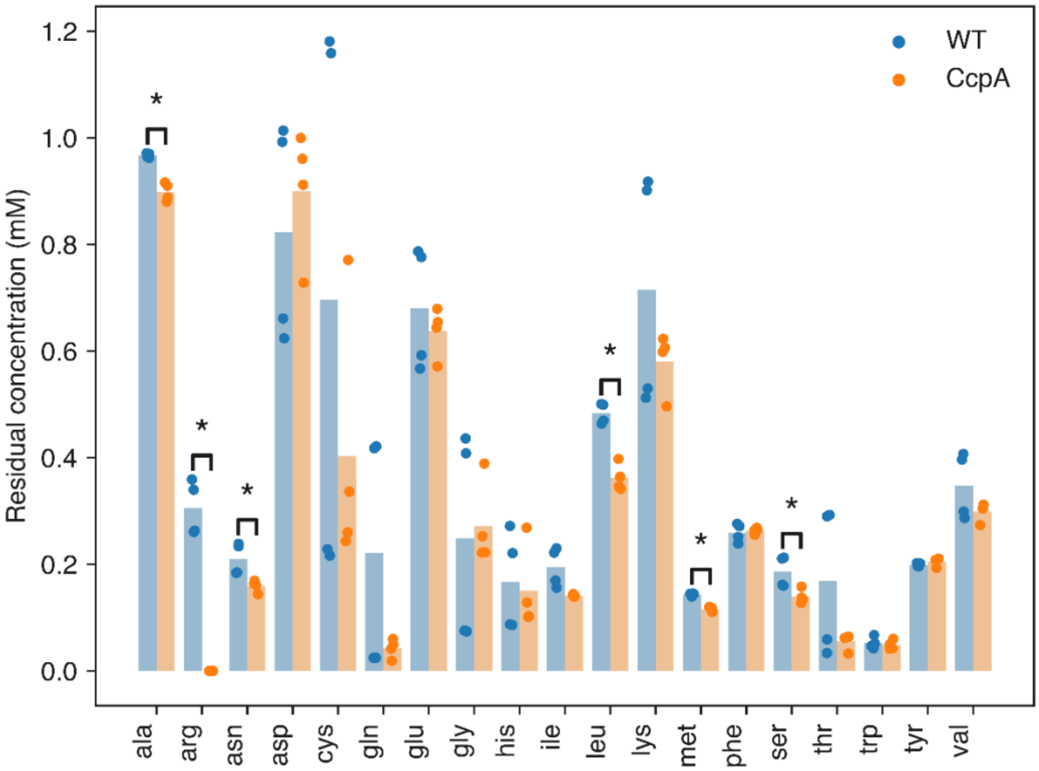


**Supplementary Fig. 5. Residual amino acid concentrations for WT *L. lactis* MG1363 and the CcpA mutant 445C1.** Arginine residual concentration became undetectable in the CcpA mutant, all others are in excess. Data points represent the individual chemostats; bars are average values; * indicates a significant difference between WT and CcpA mutant, $p < 0.05$ for independent t-test.

**Supplementary Table 1. Membrane protein fraction.** The membrane protein fraction out of total protein is calculated according to two different annotations and the combination: the sum of protein fractions of proteins that are annotated as transmembrane in UniProt and the sum of protein fractions annotated as transporter proteins in the model. Measured proteins in membrane and non-membrane were compared with a Fisher exact test, which focuses on the distribution of significantly changed proteins over the groups. The *p* values show that the chance to have a significant change in expression of a membrane protein is higher compared to random changes in all fractions.

| | WT (MG1363) | CcpA mutant (445C1) | Fisher exact p-value |
|---|---|---|---|
| Transmembrane | 0.034 | 0.033 | 0.039 |
| Transporter | 0.085 | 0.077 | 0.049 |
| Combined | 0.107 | 0.097 | 0.030 |
