## Supplementary Note - pcLactis Construction for "Proteome constraints reveal targets for improving microbial fitness in nutrient-rich environments"

### Construction of proteome-constrained model of *L. lactis*

In this document we show a detailed description of proteome-constrained model for *Lactococcus lactis* MG1363. In addition to genome-scale metabolism, the model includes many processes in gene expression: transcription, stable RNA cleavage, mRNA degradation, tRNA modification, rRNA modification, tRNA charging, ribosomal assembly, translation, protein maturation, protein assembly, enzyme formation, and protein degradation. Besides, some other reactions have been formulated for modelling purpose, including generic RNA renaming, enzyme dilution and RNA dilution reactions. We have developed a Matlab package to construct the model, which requires the COBRA toolbox[1]. The package as well as the modelling approach are only for *L. lactis* here but expected to be used in future for other prokaryotes with existing genome-scale metabolic models (M models). The construction process is divided into five steps: information collection, reformulation of M model, construction of gene expression model (E model), formulation of pseudo reactions for modelling purpose, and determination of constraints.

### 1 Information collection

#### 1.1 Coverage of E model

We determined the coverage (the number of proteins and RNA) of E model using two methods. The first method is comparative genomics analysis. We used published ME models of *Escherichia* coli[2] and *Thermotoga maritima*[3] as templates for gene orthology analysis using the EggNOG database[4] (download 15-Apr-2015 09:03). Briefly, we imported all the protein genes from E models of *E. coli* and *T. maritima* and then searched in the database the orthologous genes in *L. lactis*. As a result, we collected 108 orthologous genes in *L. lactis* compared to *E. coli* (194) and 112 compared to *T. maritima* (159). By combining these two, we obtained 134 protein genes. The second method is subsystem analysis. We used the SEED subsystem (http://pubseed.theseed.org/) to collect protein genes in the gene expression process. We searched for "Lactococcus lactis subsp. cremoris MG1363", chose the genome annotation version (416870.9), exported subsystem information, and selected all the annotated genes in the category "RNA metabolism" and "Protein metabolism" (choosing these two categories is because most of the selected genes based on the first step are located in these two categories). The subsystem analysis resulted in 187 protein genes. Combining the results from two methods, we obtained in total 218 protein genes.

Next, we manually selected protein genes and then assigned them to each process or functional machinery in the E model. As a result, we determined 157 protein genes for the gene expression process in *L. lactis*.

#### 1.2 RNA

Apart from protein genes, RNA genes are essential in gene expression process as they make mRNA and tRNA as transcription and translation machineries. We collected RNA genes from NCBI database. There are 19 rRNA and 62 tRNA genes in *L. lactis* MG1363, in total 81 RNA genes.

#### 1.3 Sequence information

We downloaded chromosome sequence information from NCBI database: https://www.ncbi.nlm.nih.gov/nuccore/NC_009004.1. All the base sequences correspond to the base sequences of the DNA coding strand, i.e., same as transcripts produced (with thymine replaced by uracil). Besides, locus ID names should be collected as there are two types of ID used in NCBI, i.e., old and new ID. We downloaded the ID relationship from BioCyc database[5] for converting ID names.

#### 1.4 Transcription unit

We collected all the transcription units (TUs) for *L. lactis* MG1363. The model produces TU as transcription product rather than transcript of a single gene. We downloaded all the TUs from BioCyc database (Lactococcus lactis, Subspecies cremoris, Strain MG1363, version 21.5).

#### 1.5 RNA modification

We assumed that transcribed tRNA and rRNA should be modified before they participate in gene expression process. We collected a great number of post-transcriptional modifications of RNA, which is a comprehensive dataset for *L. lactis*.

We determined tRNA modifications based on a published paper[6] and bioinformatics predictions[7]. The published paper experimentally identified 16 different types of modifications in 40 tRNA in *L. lactis*. Then, we used the bioinformatics analysis on the website (http://genesilico.pl/trnamodpred/) to predict the remaining modifications which cannot be identified experimentally or detected by standard mass spectrometry approaches, e.g., pseudouridine. We filtered out the resulting predictions using the term "gram positive bacteria". Subsequently, we determined proteins responsible for the modification events based on the predicted enzymes and genome annotations. For most of the modification events we can find the catalysts, but we cannot find genes responsible for C:ac4C and U:mo5U, so we assumed that these two modification events occurred spontaneously. At last, we formulated all the tRNA modification reactions as done in the ME models of *E. coli* and *T. maritima*.

Due to the fact that there are no organism-specific publications regarding rRNA modification in *L. lactis*, we determined rRNA modification events for *L. lactis* based on the information from *E. coli*. We only accounted for modifications on 16S and 23S rRNA as 5S rRNA modifications are infrequent in bacteria. We aligned rRNA in *L. lactis* and *E. coli* using ClustalW2[8] and then determined modification types and sites based on the *E. coli* rRNA modification events collected before[9]. As a result, we obtained modification events at 10 (11 in *E. coli*) positions for 16S rRNA and 24 (24 in *E. coli*) positions for 23S rRNA in *L. lactis*. Using the same method as mentioned in tRNA modification determination process, we collected genes encoding catalysts based on KO, COG or KEGG BLAST. At last, we formulated all the rRNA modification reactions as done in the ME models of *E. coli* and *T. maritima*.

#### 1.6 Protein stoichiometry

For each functional protein, we should determine whether its functional unit is a monomer or oligomer. Firstly, we downloaded all the protein sequences from the RCSB Protein Data Bank[10] (download 2018.10.30). Then, we used BLASTP to search the protein with reported protein stoichiometry information that closely related to a protein of interest. For a protein in *L. lactis*, we normally chose the identified protein with the highest bit scores, but we then chose the one with the lowest E values if highest bit scores correspond to more than one protein. When proteins had the same bit scores and E values, we chose the simplest structure from PDB. Note that we adopted the protein stoichiometry information from the biological assembly rather than the asymmetric unit as the former one is believed to be the functional form of the protein molecular (http://pdb101.rcsb.org/learn/guide-to-understanding-pdb-data/biological-assemblies). We only considered the proteins with identity >= 30% and coverage >= 80%. When no similar proteins were found, we assumed that the protein of interest was a monomer. This resulted in a comprehensive dataset of protein stoichiometry for *L. lactis*.

#### 1.7 EC number

We collected EC numbers for proteins in *L. lactis*, which can be used to retrieve $k_{cat}$ values. The EC numbers were downloaded from the UniProt[11] and KEGG[12] database.

##### 1.8 N-terminus prediction

Due to the fact that the N-terminus is usually cleaved after a peptide has been translated, we predicted for each protein whether or not the N-terminal methionine should be cleaved. This was performed using TermiNator[13] (https://bioweb.i2bc.paris-saclay.fr/terminator3/). Before running the prediction, we selected "prokaryote", "eubacteria", "intrinsic, chromosome-encoded gene", and "no LPR cleavage". Then we uploaded the sequences of all proteins and ran the prediction. We only chose the proteins with likelihood > 80%, while for the rest we assumed that there was no N-terminal methionine cleavage.

#### 2 M model reformulation

In this step, we updated the existing genome-scale metabolic model of *L. lactis*[14] and then reformulated the reactions to enable integration with E model.

##### 2.1 Reconstructing the M model

Firstly, we updated metabolic and transport reactions as well as gene-protein-reaction associations (GPRs). The workflow is shown in Fig. 1. We used MetaDraft version 0.7.2 (doi:10.5281/zenodo.2398336), a tool that reconstructs genome-scale metabolic models based on previous manually curated ones by gene homology. As source models we used the existing *L. lactis* MG1363 model[14], and the default models in the MetaDraft database with the following importance ranking: the Lactic Acid Bacteria (LAB) models (*L. lactis* model[15]), *Lactobacillus plantarum* model[16], then the remaining BiGG models[17] that were present in the default MetaDraft database (iJO1366, iAF692, iYO844, iHN637, iAF987, iIT341, iYL1228, iJN746, iSB619, iMM904, iS_1188, STM_v1_0). We manually checked all the reactions, whether the reaction and annotated genes were already present in the existing metabolic model and whether the annotated function was seen for *L. lactis* in databases. Each newly added reaction was checked within UniProt, string gene database and Artemis, to find an indication for the presence of this gene-function combination in *L. lactis* MG1363. We then conducted a search for transporter genes in *L. lactis* with the TransportDB to improve GPRs. Furthermore, we converted all metabolite and reaction IDs to be compatible with the BiGG database for collecting (Miriam) annotations and for reaction mapping and visualization purposes. This was done with the assistance of a semi-automated algorithm as described in[18].

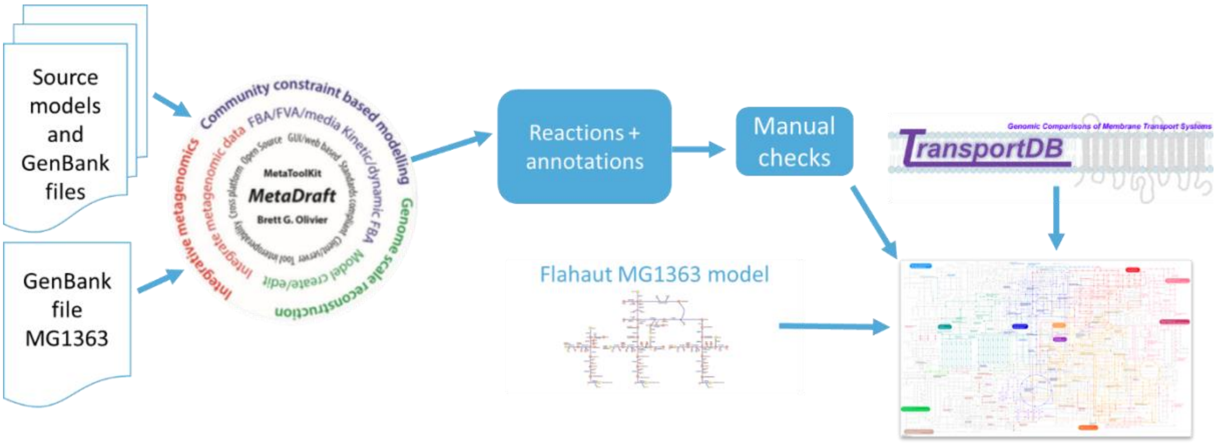

**Fig. 1. Overview of M model update.**

At last, we adjusted the biomass equation of the metabolic model for practical reasons: all tRNA charging reactions were removed from the model, because they should be present in the E model. We therefore also adjusted the biomass protein reaction to include the single amino acid in the correct ratio instead of the tRNA loaded with amino acid.

#### 2.2 Changing unknown genes

There are some unknown genes in the M model because of missing annotations. The unknown genes cannot be ignored, especially for the case that a complex has several genes but most of them are unknown, removing the unknown genes will thereby decrease the usage of materials and energy for producing the complex. Therefore, we made rules to cope with unknown genes. For the case "(unknown or geneA)", we removed the unknown gene. For the case "(unknown and geneA)", we assumed that the complex was made up of two geneA protein, i.e., the unknown one is regarded as geneA.

#### 2.3 Adding dummy GPR

There are also a lot of reactions without GPRs due to missing information in genome annotations. We assigned a "dummy" monomer to represent the enzyme for these reactions to eliminate the bias toward using these reactions without energy and material cost. We did not assign the "dummy" GPR for spontaneous reactions, e.g., "L glutamate 5 semialdehyde dehydratase". The length of the "dummy" enzyme was assumed to be a monomer with 636 amino acids, the composition was assumed to be the same as that in biomass composition, and the $k_{cat}$ value of the enzyme was assumed to be the median among other experimentally determined values.

The number of amino acids of the dummy GPR was determined as follow. Firstly, the median molecular weight of metabolic catalysts was calculated, i.e., 71503 g/mol. Then, this value was divided by the average amino acid molecular weight of 113 g/mol to obtain the number of amino acids.

#### 2.4 Splitting isozymes and reversible reactions

There are several types of GPRs in the M model, e.g., “geneA”, “(geneA and geneB)”, “(geneA or geneB)”, “(geneA and (geneB or geneC))”, “((geneA and geneB) or (geneC and geneD))”, and so on. We split each reaction with isozymes, i.e., with “or” in its GPR, into multiple reactions catalysed by different isozymes.

We also split reversible reactions into forward and reverse reactions, which was done only for enzymatic reactions.

In the following part, we use acetate kinase reaction as an example to show how a metabolic reaction was split into multiple reactions in the model.

In the M model, it is a reversible reaction catalysed by two isozymes:

- **Reaction ID:** R_ACKr
- **Reaction Formula:** M_atp_c + M_ac_c <-> M_adp_c + M_actp_c
- **Catalyst:** (llmg_2288 or llmg_2289)

Subsequently, we adjusted the reaction ID by adding “M” to mark it as metabolic reaction, and then split it into forward and reverse reactions catalysed by different isozymes. As a result, we obtained four unique reactions below:

- **Reaction ID:** R_M_ACKr_1_fwd
- **Reaction Formula:** M_atp_c + M_ac_c -> M_adp_c + M_actp_c
- **Catalyst:** M_ACKr_1_fwd_Enzyme_c

- **Reaction ID:** R_M_ACKr_1_rvs
- **Reaction Formula:** M_adp_c + M_actp_c -> M_atp_c + M_ac_c
- **Catalyst:** M_ACKr_1_rvs_Enzyme_c

- **Reaction ID:** R_M_ACKr_2_fwd
- **Reaction Formula:** M_atp_c + M_ac_c -> M_adp_c + M_actp_c
- **Catalyst:** M_ACKr_2_fwd_Enzyme_c

- **Reaction ID:** R_M_ACKr_2_rvs
- **Reaction Formula:** M_adp_c + M_actp_c -> M_atp_c + M_ac_c
- **Catalyst:** M_ACKr_2_rvs_Enzyme_c

In the reaction ID we used the number “1” and “2” (and “3” or more for the case that has three or more isozymes) to distinguish reactions catalysed by different isozymes and “fwd” and “rvs” to represent forward and reverse direction, respectively. It should be noted that we used a new term, e.g., “M_ACKr_1_fwd_Enzyme_c”, which is derived from the reaction ID, to represent catalyst rather than the original gene name in the M model. This enables each enzymatic reaction to have its own and unique catalyst ID, and thereby makes it possible to perform coupling constraints. Therefore, we formulated an extra reaction representing the formation of catalyst for each reaction. We moved such reaction in a new subprocess named “Enzyme formation”. For the four reactions above, we formulated reactions for catalyst formation:

- **Reaction ID:** R_M_ACKr_1_fwd_Enzyme
- **Reaction Formula:** llmg_2288_2mer_c -> M_ACKr_1_fwd_Enzyme_c
- **Catalyst:** -

- **Reaction ID:** R_M_ACKr_1_rvs_Enzyme
- **Reaction Formula:** llmg_2288_2mer_c -> M_ACKr_1_rvs_Enzyme_c
- **Catalyst:** -

- **Reaction ID:** R_M_ACKr_2_fwd_Enzyme
- **Reaction Formula:** llmg_2289_2mer_c -> M_ACKr_2_fwd_Enzyme_c
- **Catalyst:** -

- **Reaction ID:** R_M_ACKr_2_rvs_Enzyme
- **Reaction Formula:** llmg_2289_2mer_c -> M_ACKr_2_rvs_Enzyme_c
- **Catalyst:** -

In addition, we made dilution reactions for all the catalysts in the model, which belong to a subprocess named “Enzyme dilution”. For the example here, we also had four dilution reactions:

- **Reaction ID:** R_dilution_M_ACKr_1_fwd_Enzyme
- **Reaction Formula:** M_ACKr_1_fwd_Enzyme_c ->
- **Catalyst:** -

- **Reaction ID:** R_dilution_M_ACKr_1_rvs_Enzyme
- **Reaction Formula:** M_ACKr_1_rvs_Enzyme_c ->
- **Catalyst:** -

- **Reaction ID:** R_ dilution_M_ACKr_2_fwd_Enzyme
- **Reaction Formula:** M_ACKr_2_fwd_Enzyme_c ->
- **Catalyst:** -

- **Reaction ID:** R_ dilution_M_ACKr_2_rvs_Enzyme
- **Reaction Formula:** M_ACKr_2_rvs_Enzyme_c ->
- **Catalyst:** -

### 3 E model construction

E model accounts for synthesis of proteins used in both M and E model. We formulated reactions for transcription, stable RNA cleavage, mRNA degradation, rRNA modification, tRNA modification, ribosomal assembly, tRNA charging, translation, protein maturation, and protein assembly. Besides, we formulated generic RNA renaming reactions for modelling purpose.

#### 3.1 Transcription

Transcription is the first step of gene expression, which, using DNA as a template, consumes nucleotides and energy to produce TUs. The model only accounts for the TUs containing the protein and RNA genes involved in metabolism and gene expression. We divided transcription process into two steps, i.e., 1) transcription initiation, and 2) transcription elongation and termination.

##### 3.1.1 Transcription initiation

Transcription initiation is catalysed by RNA polymerase together with one or more sigma factors. In *L. lactis*, the core RNA polymerase consists of two alpha subunits (llmg_2354), one beta subunit (llmg_1982), one beta prime subunit (llmg_1981), one omega subunit (llmg_2154), and one delta subunit (llmg_0608). Besides, there is only one sigma factor in *L. lactis* according to genome annotation, i.e., RNA polymerase sigma factor RpoD (llmg_0521). We formulated a reaction representing the formation of RNA polymerase with sigma factor:

- **Reaction ID:** R_RNAP_sf_Enzyme

- **Reaction Formula:** llmg_2354_2mer_c + llmg_1982_Monomer_c + llmg_1981_assumed_Monomer_c + llmg_2154_assumed_Monomer_c + llmg_0608_assumed_Monomer_c + llmg_0521_Monomer_c -> RNAP_sf_Enzyme_c
- **Catalyst:** -

During the transcription initiation step, we assumed that the first 16 nucleotides were transcribed and 15 diphosphate (ppi) were released. We use the TU "TU1G1R-100" as an example to show transcription initiation below:

- **Reaction ID:** R_TU1G1R_100_Initiation
- **Reaction Formula:** 6 M_atp_c + M_gtp_c + 7 M_utp_c + 2 M_ctp_c -> 15 M_ppi_c + TU1G1R_100_Initiated_c
- **Catalyst:** RNAP_sf_Enzyme_c

##### 3.1.2 Transcription elongation and termination

We lumped transcription elongation and termination to decrease the total number of reactions. From the KEGG database (Transcription machinery [BR: llm03021]), we collected transcription elongation factor NusA (llmg_1796), transcription termination protein NusB (llmg_1878), transcription antitermination protein NusG (llmg_2388), and transcription elongation factor GreA (llmg_0610). Besides, we manually added transcription-repair coupling factor Mfd (llmg_0013). We cannot find Rho termination factor in *L. lactis* genome, so all the transcription reactions are Rho independent. We formulated the formation of transcription complex, which consists of all the factors mentioned above:

- **Reaction ID:** R_Transcription_Complex_Enzyme
- **Reaction Formula:** llmg_1796_2mer_c + llmg_1878_assumed_Monomer_c + llmg_2388_Monomer_c + llmg_0610_Monomer_c + llmg_0013_Monomer_c -> Transcription_Complex_Enzyme_c
- **Catalyst:** -

Catalysed by the transcription complex, the initiated TU will be elongated and terminated in this step (still using "TU1G1R_100" as an example):

- **Reaction ID:** R_TU1G1R_100
- **Reaction Formula:** 304 M_atp_c + 215 M_gtp_c + 276 M_utp_c + 191 M_ctp_c + TU1G1R_100_Initiated_c -> 986 M_ppi_c + TU1G1R_100_c
- **Catalyst:** Transcription_Complex_Enzyme_c

#### 3.2 Stable RNA cleavage

There are two types of TUs produced by transcription process, one type is the TU exclusively containing protein genes while the other is the TU containing RNA genes. The former will be translated directly, but the latter should be cleaved before entering other subprocesses as RNA genes will not be translated. Therefore, we added stable RNA cleavage process for the TUs that contain RNA genes.

Ribonuclease is responsible for stable RNA cleavage and there are many types of ribonucleases in *L. lactis* according to genome annotation and the KEGG database. They are RNase III (llmg_1753), mini-RNase III (llmg_2039), RNase P (llmg_0142), RNase HII (llmg_1176), RNase HIII (llmg_2549), RNase M5 (llmg_1883), RNase Z (llmg_0412), RNase Z (llmg_0604), RNase J1 (llmg_0302), RNase BN (llmg_0578), RNase J2 (llmg_0876), RNase R (llmg_1304), and RNase R (llmg_1586). However, we only assumed RNase III (llmg_1753) to catalyse RNA cleavage reactions as we cannot find any supporting evidence. So, we formulated the formation of RNase III:

- **Reaction ID:** R_RNase_Enzyme
- **Reaction Formula:** llmg_1753_2mer_c -> RNase_Enzyme_c
- **Catalyst:** -

For the case that a TU contains only one RNA gene, we formulated a spontaneous reaction:

- **Reaction ID:** R_TU1G1R_1030_cleavage
- **Reaction Formula:** TU1G1R_1030_c -> llmg_tRNA_36_unmodified_c
- **Catalyst:** -

For the case that a TU contains more than one RNA gene, RNase III should be used. It should be noted that one $H_2O$ should be used per time of cleavage for one TU, and one proton is produced. Besides, sub-TU could be produced in this process, which is the sequence between two RNA genes in the TU. The sub-TU can be either degraded directly or translated if it contains protein genes. Below is an example:

- **Reaction ID:** R_TU1G1R_3106_cleavage
- **Reaction Formula:** 10 M_h2o_c + TU1G1R_3106_c -> 10 M_h_c + llmg_rRNA_60a_unmodified_c + llmg_rRNA_7_unmodified_c + llmg_rRNA_7a_unmodified_c + llmg_tRNA_62_unmodified_c + llmg_tRNA_61_unmodified_c + llmg_tRNA_60_unmodified_c + TU1G1R_3106_sub_1_c + TU1G1R_3106_sub_2_c + TU1G1R_3106_sub_3_c + TU1G1R_3106_sub_4_c + TU1G1R_3106_sub_5_c
- **Catalyst:** RNase_Enzyme_c

#### 3.3 mRNA degradation

We formulated mRNA degradation reactions for all the TUs produced in transcription process and sub-TUs produced in stable RNA cleavage process. The catalysts for this process are degradosome and oligoribonuclease. In *L. lactis*, the degradosome seems to be RNA degradosome type D according to the KEGG database, which is similar to *Bacillus subtilis*. The type D degradosome is made up of RNaseY (llmg_2156), CshA (llmg_0369), RNaseJ1 (llmg_0302), RNaseJ2 (llmg_0876), Enolase (llmg_0617), PNPase (llmg_2044), and PfkA (llmg_1118). Thereby, we formulated the formation of the mRNA degradation complex by integrating degradosome and oligoribonuclease Orn (llmg_1825):

- **Reaction ID:** R_mRNA_Degradation_Complex_Enzyme
- **Reaction Formula:** llmg_0617_Monomer_c + llmg_1118_4mer_c + llmg_0302_4mer_c + llmg_0369_assumed_Monomer_c + llmg_0876_4mer_c + llmg_1825_Monomer_c + llmg_2044_3mer_c + llmg_2156_assumed_Monomer_c -> mRNA_Degradation_Complex_Enzyme_c
- **Catalyst:** -

The mRNA degradation process is energy-consuming, and we assumed that one ATP was needed for every four nucleotides degraded as done for modelling *E. coli* and *T. maritima*. Nucleotide within TU was hydrolysed into nucleoside monophosphate except the one at the first position, which was returned as nucleoside triphosphate. We assumed that the degradation of one molecule of TU or sub-TU was catalysed by only one molecule of mRNA degradation complex regardless of the number of nucleotides. Below is an example:

- **Reaction ID:** R_TU1G1R_1034_degradation
- **Reaction Formula:** 324 M_atp_c + 1619 M_h2o_c + TU1G1R_1034_c -> 418 M_amp_c + 1619 M_h_c + M_gtp_c + 324 M_pi_c + 324 M_adp_c + 385 M_ump_c + 213 M_cmp_c + 279 M_gmp_c
- **Catalyst:** mRNA_Degradation_Complex_Enzyme_c

#### 3.4 tRNA modification

RNA modification is a process that some bases of an RNA molecule are changed. We modelled this process by changing one base at a time. There are three cases in the model, i.e., 1) modification without any metabolites consumed or produced, 2) modification without any catalysts, and 3) modification with metabolites and catalysts involved. We show the examples below for the three cases:

1) modification without any metabolites consumed or produced

- **Reaction ID:** R_tRNA_modification_llmg_tRNA_27_modified_1
- **Reaction Formula:** llmg_tRNA_27_unmodified_c -> llmg_tRNA_27_modified_1_c

- **Catalyst:** RNAmod_llmg_0395_Enzyme_c

2) modification without any catalysts

- **Reaction ID:** R_tRNA_modification_llmg_tRNA_27_modified_3
- **Reaction Formula:** M_accoa_c + llmg_tRNA_27_modified_2_c -> M_coa_c + llmg_tRNA_27_modified_3_c
- **Catalyst:** -

3) modification with metabolites and catalysts involved

- **Reaction ID:** R_tRNA_modification_llmg_tRNA_27_modified_4
- **Reaction Formula:** M_amet_c + llmg_tRNA_27_modified_3_c -> M_h_c + M_ahcys_c + llmg_tRNA_27_modified_4_c
- **Catalyst:** RNAmod_llmg_2424_Enzyme_c

The formation reactions of all the catalysts were also formulated in this stage and stored in "Enzyme formation" subprocess.

#### 3.5 Generic RNA renaming

We formulated generic RNA renaming reactions for tRNA and rRNA to save the number of reactions. This was done for tRNA after having been modified as different tRNA even sharing the same codon may show different sequences and thereby different modification events, but for rRNA before being modified as the same type of rRNA have identical sequences and thereby the same modification events. We show the renaming of tRNA-Val-TAC, which can be made by three genes in *L. lactis*, as an example below:

- **Reaction ID:** R_Generic_RNA_llmg_tRNA_19_tRNA_Val_GUA_to_tRNA_Val_GUA
- **Reaction Formula:** llmg_tRNA_19_tRNA_Val_GUA_c -> tRNA_Val_GUA_c
- **Catalyst:** -

- **Reaction ID:** R_Generic_RNA_llmg_tRNA_47_tRNA_Val_GUA_to_tRNA_Val_GUA
- **Reaction Formula:** llmg_tRNA_47_tRNA_Val_GUA_c -> tRNA_Val_GUA_c
- **Catalyst:** -

- **Reaction ID:** R_Generic_RNA_llmg_tRNA_58_tRNA_Val_GUA_to_tRNA_Val_GUA
- **Reaction Formula:** llmg_tRNA_58_tRNA_Val_GUA_c -> tRNA_Val_GUA_c
- **Catalyst:** -

We show the renaming of 16S-rRNA, which can be made by six genes in *L. lactis*, as an example below:

- **Reaction ID:** R_Generic_RNA_llmg_rRNA_1_unmodified_to_rRNA_16S_unmodified
- **Reaction Formula:** llmg_rRNA_1_unmodified_c -> rRNA_16S_unmodified_c
- **Catalyst:** -

- **Reaction ID:** R_Generic_RNA_llmg_rRNA_4a_unmodified_to_rRNA_16S_unmodified
- **Reaction Formula:** llmg_rRNA_4a_unmodified_c -> rRNA_16S_unmodified_c
- **Catalyst:** -

- **Reaction ID:** R_Generic_RNA_llmg_rRNA_5a_unmodified_to_rRNA_16S_unmodified
- **Reaction Formula:** llmg_rRNA_5a_unmodified_c -> rRNA_16S_unmodified_c
- **Catalyst:** -

- **Reaction ID:** R_Generic_RNA_llmg_rRNA_6b_unmodified_to_rRNA_16S_unmodified
- **Reaction Formula:** llmg_rRNA_6b_unmodified_c -> rRNA_16S_unmodified_c
- **Catalyst:** -

- **Reaction ID:** R_Generic_RNA_llmg_rRNA_7a_unmodified_to_rRNA_16S_unmodified
- **Reaction Formula:** llmg_rRNA_7a_unmodified_c -> rRNA_16S_unmodified_c
- **Catalyst:** -

- **Reaction ID:** R_Generic_RNA_llmg_rRNA_48a_unmodified_to_rRNA_16S_unmodified
- **Reaction Formula:** llmg_rRNA_48a_unmodified_c -> rRNA_16S_unmodified_c
- **Catalyst:** -

#### 3.6 rRNA modification

The formulation of rRNA modification reaction is similar to tRNA modification. We formulated modification reactions only for 16S rRNA and 23S rRNA as modifications to 5S rRNA seem to be infrequent in bacteria. We predicted that in *L. lactis* there were ten modification sites in 16S rRNA and 24 modification events in 23S rRNA. Therefore, we formulated ten reactions for 16S rRNA modification and 24 reactions for 23S rRNA modification. We show the first and the last reaction of 16S rRNA modification below.

The first reaction:

- **Reaction ID:** R_rRNA_modification_rRNA_16S_modified_1
- **Reaction Formula:** rRNA_16S_unmodified_c -> rRNA_16S_modified_1_c
- **Catalyst:** RNAmod_llmg_2518_Enzyme_c

The last reaction:

- **Reaction ID:** R_rRNA_modification_rRNA_16S
- **Reaction Formula:** 2 M_amet_c + rRNA_16S_modified_9_c -> 2 M_h_c + 2 M_ahcys_c + rRNA_16S_c
- **Catalyst:** RNAmod_llmg_1882_Enzyme_c

#### 3.7 Ribosomal assembly

We divided ribosomal assembly into two steps, i.e., ribosome protein formation and assembly of protein and rRNA. Firstly, we formulated formation of each ribosome protein subunit, which is spontaneous in the model. It should be noted that in *L. lactis* ribosome protein subunit L33 can be coded by three genes, i.e., "llmg_0098", "llmg_0632", and "llmg_2390". We found in the proteome of *L. lactis* Il1403 that only the protein of "L0096" (llmg_0098) was detected with its abundance in top 5% of the total detected proteome[19]. Therefore, we assumed that only "llmg_0098" was used for the production of L33. Besides, the coefficient of L7/L12 (llmg_1208) was assumed to be four according to the fact that L12 presents in four copies in prokaryotic ribosomes[20].

The formation of 30S ribosomal protein was formulated by adding all the small subunits:

- **Reaction ID:** R_ribosome_30S_protein
- **Reaction Formula:** llmg_0251_Monomer_c + llmg_0296_Monomer_c + llmg_0932_Monomer_c + llmg_1724_assumed_Monomer_c + llmg_1921_Monomer_c + llmg_2078_Monomer_c + llmg_2355_Monomer_c + llmg_2356_Monomer_c + llmg_2364_Monomer_c + llmg_2367_Monomer_c + llmg_2370_Monomer_c + llmg_2374_Monomer_c + llmg_2377_Monomer_c + llmg_2379_Monomer_c + llmg_2384_Monomer_c + llmg_2430_Monomer_c + llmg_2473_Monomer_c + llmg_2475_Monomer_c + llmg_2545_Monomer_c + llmg_2557_Monomer_c + llmg_2558_Monomer_c -> ribosome_30S_protein_c
- **Catalyst:** -

The formation of 50S ribosomal protein was formulated by adding all the large subunits:

- **Reaction ID:** R_ribosome_50S_protein

- **Reaction Formula:** llmg_0098_Monomer_c + llmg_0099_Monomer_c + llmg_0145_Monomer_c + llmg_0204_Monomer_c + llmg_0906_Monomer_c + llmg_1207_Monomer_c + 4 llmg_1208_Monomer_c + llmg_1491_Monomer_c + llmg_1493_Monomer_c + llmg_1671_Monomer_c + llmg_1815_Monomer_c + llmg_2029_Monomer_c + llmg_2030_Monomer_c + llmg_2276_Monomer_c + llmg_2277_Monomer_c + llmg_2353_Monomer_c + llmg_2357_Monomer_c + llmg_2362_Monomer_c + llmg_2363_Monomer_c + llmg_2365_Monomer_c + llmg_2366_Monomer_c + llmg_2371_Monomer_c + llmg_2372_Monomer_c + llmg_2373_Monomer_c + llmg_2375_Monomer_c + llmg_2376_Monomer_c + llmg_2378_Monomer_c + llmg_2380_Monomer_c + llmg_2381_Monomer_c + llmg_2382_Monomer_c + llmg_2383_Monomer_c + llmg_2546_Monomer_c -> ribosome_50S_protein_c
- **Catalyst:** -

The 70S ribosome is made up of 30S and 50S ribosome, each of which is made up of ribosomal protein and rRNA. For 30S ribosome, it is made up of 16S rRNA and 30S ribosomal protein. We assumed that the formation of 30S ribosome was catalysed by a complex consisting of GTP-binding protein Era (llmg_0371), Ribosome-binding factor A (RbfA) (llmg_1791), and 16S rRNA processing protein RimM (llmg_0936). Besides, the formation of one molecule of 30S ribosome needs one molecule of GTP as energy.

Below is the formation of 30S ribosome catalyst complex:

- **Reaction ID:** R_Ribosome_30S_Enzyme
- **Reaction Formula:** llmg_0371_Monomer_c + llmg_1791_Monomer_c + llmg_0936_2mer_c -> Ribosome_30S_Enzyme_c
- **Catalyst:** -

Therefore, we formulated the formation of 30S ribosome below:

- **Reaction ID:** R_ribosome_30S
- **Reaction Formula:** M_gtp_c + rRNA_16S_c + ribosome_30S_protein_c -> M_pi_c + M_h_c + M_gdp_c + ribosome_30S_c
- **Catalyst:** Ribosome_30S_Enzyme_c

For 50S ribosome, it is made up of 5S rRNA, 23S rRNA, and 50S ribosomal protein. We assumed that the formation of 50S ribosome was catalysed by trigger factor (llmg_0519) without energy cost.

Below is the formation of 50S ribosome catalyst:

- **Reaction ID:** R_Ribosome_50S_Enzyme
- **Reaction Formula:** llmg_0519_2mer_c -> Ribosome_50S_Enzyme_c
- **Catalyst:** -

Therefore, we formulated the formation of 50S ribosome below:

- **Reaction ID:** R_ribosome_50S
- **Reaction Formula:** rRNA_5S_unmodified_c + rRNA_23S_c + ribosome_50S_protein_c -> ribosome_50S_c
- **Catalyst:** Ribosome_50S_Enzyme_c

At last, we formulated the formation of 70S ribosome, which is spontaneous in the model:

- **Reaction ID:** R_ribosome_70S
- **Reaction Formula:** ribosome_30S_c + ribosome_50S_c -> ribosome_70S_c
- **Catalyst:** -

#### 3.8 tRNA charging

We modelled tRNA charging process for 1) charging tRNA with amino acids and 2) generating charged codon readers for the codons that do not have corresponding tRNA.

In *L. lactis*, there are 35 types of tRNA with various anticodons. It should be noted that both tRNA-Met-CAU and tRNA-fMet-CAU have the anticodon “CAU” but they function differently, i.e., the latter transfers Met for the first position while the former for the others. Firstly, we formulated tRNA charging reactions for all the 35 types of tRNA, and the reactions were based on the KEGG database. Generally, tRNA charging is a multiple-step process and catalysed by tRNA synthetase. We lumped the multiple steps in the model and used one enzymatic reaction to represent this process. It should be noted that we did not include tRNA as a substrate in the reaction as otherwise it will be returned as a product in translation process and thereby complicate the network. Instead, we coupled generic tRNA renaming reactions with tRNA charging reactions to account for the need of tRNA when solving the model. Below is an example for charging tRNA with the anticodon “GCA”:

- **Reaction ID:** R_A_charging_tRNA_Ala_GCA
- **Reaction Formula:** M_ala__L_c + M_atp_c + M_h2o_c -> A_charged_in_tRNA_Ala_GCA_c + M_amp_c + M_ppi_c + M_h_c
- **Catalyst:** tRNA_Synthetase_A_GCA_Enzyme_c

When charging tRNA-fMet, which is responsible for transferring methionine for the first codon, we integrated formyltransferase reaction into the charging reaction. Thereby, the integrated reaction was assumed to be catalysed by a complex consisting of methionyl-tRNA synthetase and methionyl-tRNA formyltransferase. Below is an example:

- **Reaction ID:** R_fM_charging_tRNA_fMet_AUG
- **Reaction Formula:** M_atp_c + 2 M_h2o_c + M_10fthf_c + M_met__L_c -> M_amp_c + M_ppi_c + 2 M_h_c + M_thf_c + fM_charged_in_tRNA_fMet_AUG_c
- **Catalyst:** tRNA_Synthetase_Complex_fM_AUG_Enzyme_c

Notably, there is no glutamine-tRNA synthetase gene in *L. lactis* genome, so tRNA-gln is charged in a different way. Briefly, glutamate is bound to tRNA-Gln by glutamyl-tRNA synthetase, and then amidotransferase converts the bound glutamate to glutamine. We lumped the two steps in the model and obtained one reaction catalysed by a complex consisting of glutamyl-tRNA synthetase and amidotransferase. Below is an example:

- **Reaction ID:** R_Q_charging_tRNA_Gln_CAA
- **Reaction Formula:** 2 M_atp_c + M_h2o_c + M_gln__L_c -> M_amp_c + M_ppi_c + M_h_c + M_pi_c + Q_charged_in_tRNA_Gln_CAA_c + M_adp_c
- **Catalyst:** tRNA_Synthetase_Complex_Q_CAA_Enzyme_c

There are $4^3$ = 64 possible codons, so there should be 61 (excluding three stop codons) types of codon readers (i.e., tRNA) if each one follows base pair rules. However, there are 35 types of tRNA in *L. lactis* which only correspond to 34 anticodons, so we generated codon readers for the remaining codons (61 – 34 = 27). The biological explanation behind this is the Wobble hypothesis, i.e., the 5’ base on the anticodon (or the 3’ base on the codon) could have flexible choices of paired base. Therefore, we formulated reaction for the formation of codon readers for the remaining 27 codons. Considering the fact that a codon could be read by more than one tRNA when following the Wobble hypothesis, e.g., tRNA with anticodon “GCC” and “UCC” can both read the codon “GGU”, we assumed that only the tRNA with the lowest cost in its production was used. For simplification, we formulated a spontaneous reaction to obtain a charged codon reader directly from a charged tRNA. We show below tRNA-Gly as an example. There are four codons that can be translated to glycine, but only two codon readers in *L. lactis*, i.e., tRNA with anticodon “UCC” and that with “GCC”. Following base pair rules, they can read the codon “GGA” and “GGC”, respectively, and the remaining codons are “GGG” and “GGU”, which can be read by both of them following Wobble hypothesis. However, we only used the tRNA with anticodon "UCC" to produce the codon readers for both the codon "GGG" and "GGU" as it is shorter in length than the tRNA with anticodon "GCC". Below are reactions:

- **Reaction ID:** R_G_charged_in_tRNA_Gly_GGG
- **Reaction Formula:** G_charged_in_tRNA_Gly_GGA_c -> G_charged_in_tRNA_Gly_GGG_c
- **Catalyst:** -

- **Reaction ID:** R_G_charged_in_tRNA_Gly_GGU
- **Reaction Formula:** G_charged_in_tRNA_Gly_GGA_c -> G_charged_in_tRNA_Gly_GGU_c
- **Catalyst:** -

#### 3.9 Translation

We divided translation process into three steps in the model, i.e., 1) translation initiation, 2) translation elongation, and 3) translation termination.

##### 3.9.1 Translation initiation

Translation initiation represents the process that 70S ribosome and the first methionine bind a TU. We assumed that an intact TU or sub-TU was translated without being divided into transcripts of single genes. Therefore, translation of any gene in a TU needs the intact TU as a template, but translation initiation starts from the start codon of the gene of interest. Besides, we assumed that 70S ribosome was used for coupling constraints only in translation elongation step although it should be used during the entire translation process. In the model, translation initiation is catalysed by a complex, which is made up of translation initiation factor 1 (llmg_2358), 2 (llmg_1792), and 3 (llmg_2031).

- **Reaction ID:** R_Translation_Initiation_Complex_Enzyme
- **Reaction Formula:** llmg_2358_4mer_c + llmg_1792_assumed_Monomer_c + llmg_2031_assumed_Monomer_c -> Translation_Initiation_Complex_Enzyme_c
- **Catalyst:** -

It should be noted that we did not include TU as a substrate in the translation initiation reaction. Instead, we coupled TU or Sub-TU production reactions with translation initiation reactions to account for the need of mRNA in simulations. We use the gene "llmg_0145" as an example to show translation initiation reaction below:

- **Reaction ID:** R_translation_initiation_TU1G1R_202_llmg_0145
- **Reaction Formula:** M_h2o_c + M_gtp_c + fM_charged_in_tRNA_fMet_AUG_c -> M_h_c + M_pi_c + M_gdp_c + TU1G1R_202_llmg_0145_initiated_c
- **Catalyst:** Translation_Initiation_Complex_Enzyme_c

##### 3.9.2 Translation elongation

Translation elongation represents the process that amino acids are added till the ribosome reaches a stop codon. There are two steps in this process: binding of a charged tRNA to the ribosome and translocation of the charged tRNA. The former is catalysed by EF-Tu (llmg_2050) together with EF-Ts (llmg_2429), and the latter by EF-G (llmg_2556). We formulated reactions representing the binding and translocation of tRNA, resulting in a pool of amino acids charged to bound and translocated tRNA, from which any amino acids can be used directly for translation elongation. In other words, we only changed the state of amino acids instead of adding tRNA to the reactions.

Firstly, we formulated reactions representing the binding of a charged tRNA to the ribosome, catalysed by a complex consisting of EF-Tu and EF-Ts.

- **Reaction ID:** R_EF_Tu_EF_Ts_Complex_Enzyme
- **Reaction Formula:** llmg_2050_Monomer_c + llmg_2429_Monomer_c -> EF_Tu_EF_Ts_Complex_Enzyme_c
- **Catalyst:** -

This was done for all the charged tRNA, but we only show "tRNA_Ala-GCA_charged" as an example below:

- **Reaction ID:** R_Activate_A_charged_in_tRNA_Ala_GCA
- **Reaction Formula:** M_h2o_c + A_charged_in_tRNA_Ala_GCA_c + M_gtp_c -> M_h_c + A_charged_in_tRNA_Ala_GCA_activated_c + M_pi_c + M_gdp_c
- **Catalyst:** EF_Tu_EF_Ts_Complex_Enzyme_c

Secondly, we formulated reactions representing the translocation of the bound tRNA (i.e., activated tRNA in the model), catalysed by EF-G.

- **Reaction ID:** R_EF_G_Enzyme
- **Reaction Formula:** llmg_2556_Monomer_c -> EF_G_Enzyme_c
- **Catalyst:** -

The activated tRNA is then converted to an elongated state (representing translocation), and the product can be directly used in translation elongation process:

- **Reaction ID:** R_Elongate_A_charged_in_tRNA_Ala_GCA

- **Reaction Formula:** M_h2o_c + M_gtp_c + A_charged_in_tRNA_Ala_GCA_activated_c -> M_h_c + M_pi_c + M_gdp_c + A_charged_in_tRNA_Ala_GCA_elongated_c
- **Catalyst:** EF_G_Enzyme_c

As a result, we generated a pool of tRNA prepared for adding amino acids in the model. The pool contains 61 charged tRNA which can read all the possible codons in nucleotide sequences. Besides, all the tRNA in the pool are already in a "bound" and "translocated" state, meaning that no catalyst or energy is needed when using them to add amino acids. Then we formulated reactions to add the remaining amino acids based on nucleotide sequences. The reaction uses the initiated product produced in translation initiation process and produces an elongated product. Note that 70S ribosome is used in this step as catalyst, and there is no tRNA involved in the reaction. We still use the gene "llmg_0145" as an example below:

- **Reaction ID:** R_translation_elongation_TU1G1R_202_llmg_0145
- **Reaction Formula:** 2 A_charged_in_tRNA_Ala_GCA_elongated_c + 2 A_charged_in_tRNA_Ala_GCU_elongated_c + F_charged_in_tRNA_Phe_UUC_elongated_c + 3 G_charged_in_tRNA_Gly_GGA_elongated_c + 2 H_charged_in_tRNA_His_CAC_elongated_c + 5 K_charged_in_tRNA_Lys_AAA_elongated_c + K_charged_in_tRNA_Lys_AAG_elongated_c + 2 L_charged_in_tRNA_Leu_CUU_elongated_c + M_charged_in_tRNA_Met_AUG_elongated_c + N_charged_in_tRNA_Asn_AAC_elongated_c + P_charged_in_tRNA_Pro_CCA_elongated_c + Q_charged_in_tRNA_Gln_CAA_elongated_c + R_charged_in_tRNA_Arg_CGC_elongated_c + 9 R_charged_in_tRNA_Arg_CGU_elongated_c + S_charged_in_tRNA_Ser_AGC_elongated_c + S_charged_in_tRNA_Ser_UCA_elongated_c + S_charged_in_tRNA_Ser_UCU_elongated_c + T_charged_in_tRNA_Thr_ACA_elongated_c + 4 T_charged_in_tRNA_Thr_ACU_elongated_c + V_charged_in_tRNA_Val_GUC_elongated_c + V_charged_in_tRNA_Val_GUU_elongated_c + Y_charged_in_tRNA_Tyr_UAC_elongated_c + TU1G1R_202_llmg_0145_initiated_c -> 43 M_h2o_c + TU1G1R_202_llmg_0145_elongated_c
- **Catalyst:** ribosome_70S_c

##### 3.9.3 Translation termination

Translation termination represents the process that the ribosome reaches a stop codon and then peptide synthesis is terminated with the release of ribosome and peptide from mRNA. There are four key factors involved in translation termination process, i.e., RF1 (llmg_0557), RF2 (llmg_1547), RF3 (llmg_0368), and ribosome recycling factor Rrf (llmg_2284). Considering the fact that RF1 recognises the stop codons "UAA" and "UAG" while RF2 recognises "UAA" and "UGA", i.e., both RF1 and RF2 can read "UAA", we assumed that only

RF1 recognises “UAA” as it needs less cost. Then adding RF3 and Rrf, we formulated reactions for the formation of translation termination complex.

- **Reaction ID:** R_RFRrf_UAA_UAG_Enzyme
- **Reaction Formula:** llmg_0557_Monomer_c + llmg_0368_Monomer_c + llmg_2284_Monomer_c -> RFRrf_UAA_UAG_Enzyme_c
- **Catalyst:** -

- **Reaction ID:** R_RFRrf_UGA_Enzyme
- **Reaction Formula:** llmg_0368_Monomer_c + llmg_2284_Monomer_c + llmg_1547_Monomer_c -> RFRrf_UGA_Enzyme_c
- **Catalyst:** -

In the model, translation termination reaction uses the elongated product produced in translation elongation process and produces a nascent peptide with a TU returned. The reaction is catalysed by one of the two translation termination complexes, depending on the stop codon of the gene of interest. We still use the gene “llmg_0145”, which ends with the stop codon “UAA”, as an example below:

- **Reaction ID:** R_translation_termination_TU1G1R_202_llmg_0145
- **Reaction Formula:** M_h2o_c + M_gtp_c + TU1G1R_202_llmg_0145_elongated_c -> M_h_c + M_pi_c + M_gdp_c + llmg_0145_nascent_c
- **Catalyst:** RFRrf_UAA_UAG_Enzyme_c

#### 3.10 Protein maturation

We modelled protein maturation process to remove formyl-group for all the translated peptides, and methionyl-group for the case that the mature protein does not have methionine at the first position. It depends on the N-terminus prediction whether methionine should be removed.

The cleavage of the formyl-group is catalysed by peptide deformylase Def (llmg_0532). We formulated the formation of Def in the model:

- **Reaction ID:** R_Peptide_Deformylase_Enzyme
- **Reaction Formula:** llmg_0532_Monomer_c -> Peptide_Deformylase_Enzyme_c
- **Catalyst:** -

The cleavage of the methionyl-group is catalysed by methionine aminopeptidase Map (llmg_0577). We formulated the formation of Map in the model:

- **Reaction ID:** R_Methionine_Aminopeptidase_Enzyme
- **Reaction Formula:** llmg_0577_2mer_c -> Methionine_Aminopeptidase_Enzyme_c
- **Catalyst:** -

There are two cases in protein maturation process: 1) only formyl-group should be removed, and 2) both formyl-group and methionyl-group should be removed. Below, we show examples for the two cases.

1) Only formyl-group should be removed (using "llmg_0145" as an example)

- **Reaction ID:** R_deformylase_llmg_0145
- **Reaction Formula:** M_h2o_c + llmg_0145_nascent_c -> M_for_c + llmg_0145_c
- **Catalyst:** Peptide_Deformylase_Enzyme_c

2) Both formyl-group and methionyl-group should be removed (using "llmg_0073" as an example)

Firstly, the formyl-group is removed:

- **Reaction ID:** R_deformylase_llmg_0073
- **Reaction Formula:** M_h2o_c + llmg_0073_nascent_c -> M_for_c + llmg_0073_for_M_excision_c
- **Catalyst:** Peptide_Deformylase_Enzyme_c

Secondly, the methionyl-group is removed:

- **Reaction ID:** R_Met_aminopeptidase_llmg_0073
- **Reaction Formula:** M_h2o_c + llmg_0073_for_M_excision_c -> M_met__L_c + llmg_0073_c
- **Catalyst:** Methionine_Aminopeptidase_Enzyme_c

#### 3.11 Protein assembly

We modelled protein assembly process for protein folding based on protein stoichiometry information. We assumed that this process was spontaneous. The protein stoichiometry determines how many nascent peptides should be used for a functional protein. We assumed the protein without known stoichiometry information to be a monomer. Below are examples for monomer, assumed monomer, and oligomer:

- **Reaction ID:** R_llmg_0013_Monomer
- **Reaction Formula:** llmg_0013_c -> llmg_0013_Monomer_c

- **Catalyst:** -

- **Reaction ID:** R_llmg_0022_assumed_Monomer
- **Reaction Formula:** llmg_0022_c -> llmg_0022_assumed_Monomer_c
- **Catalyst:** -

- **Reaction ID:** R_llmg_0020_4mer
- **Reaction Formula:** 4 llmg_0020_c -> llmg_0020_4mer_c
- **Catalyst:** -

#### 3.12 Enzyme formation

In the model, we formulated enzyme formation reactions to represent the production of functional enzymes catalysing other reactions in metabolism and gene expression processes. We formulated such reactions for all the catalysts no matter whether it is a complex, and all of them are spontaneous. We have shown some examples above and will also show some below. If a catalyst is made up of only one folded protein, e.g., acetylglutamate kinase, the reaction is:

- **Reaction ID:** R_M_ACGK_Enzyme
- **Reaction Formula:** llmg_1755_6mer_c -> M_ACGK_Enzyme_c
- **Catalyst:** -

If a catalyst is a complex and made up of several subunits, e.g., glycogen phosphorylase, the reaction is:

- **Reaction ID:** R_M_GLCP_Enzyme
- **Reaction Formula:** llmg_1869_Monomer_c + llmg_1871_Monomer_c -> M_GLCP_Enzyme_c
- **Catalyst:** -

Note that we did not include the formation of 70S ribosome in this process. Instead, we formulated such a reaction in another process, i.e., ribosomal assembly, as it is a special catalyst made up of not only proteins but also rRNA.

#### 3.13 Protein degradation

Protein degradation process in the model includes two steps, i.e., breakdown of a catalyst (complex) into sub-proteins (subunits) and proteolysis of a protein into amino acids. Considering the fact that we formulated formation reaction even for the catalyst with only one type of protein (e.g., llmg_2556_Monomer_c -> EF_G_Enzyme_c), all the catalysts in the model should go through the two steps for degradation. We assumed that the first step was spontaneous while the second step not. In *L. lactis*, several proteases have been identified, including ClpP[21], FtsH[22], and HtrA[23]. As ClpP seems to be central in both the proteolysis of misfolded proteins and adjusting regulatory proteins[24], we assumed that protein degradation was catalysed by Clp protease complex, which is made up of one proteolytic subunit ClpP (llmg_0638) and three regulatory subunits ClpB (llmg_0986), ClpC (llmg_0615), and ClpE (llmg_0528) in *L. lactis*[25]. Regarding energy cost, it was estimated that approximately 0.25 to 1.0 ATP was needed for the recycling of each amino acid during protein degradation[26]. Therefore, we adopted the lowest value, i.e., 0.25 ATP per amino acid, for degradation of each protein.

Below is the formation of Clp protease complex:

- **Reaction ID:** R_Clp_Protease_Complex_Enzyme
- **Reaction Formula:** llmg_0638_14mer_c + llmg_0986_3mer_c + llmg_0615_10mer_c + llmg_0528_10mer_c -> Clp_Protease_Complex_Enzyme_c
- **Catalyst:** -

Here we use “tRNA_Synthetase_F_UUC_Enzyme_c” as an example to show two steps of protein degradation.

The first step is the breakdown of the complex:

- **Reaction ID:** R_tRNA_Synthetase_F_UUC_Enzyme_degradation
- **Reaction Formula:** tRNA_Synthetase_F_UUC_Enzyme_c -> llmg_2195_2mer_degradation_c + llmg_2196_2mer_degradation_c
- **Catalyst:** -

The second step is proteolysis of sub-proteins into amino acids:

- **Reaction ID:** R_llmg_2195_2mer_degradation
- **Reaction Formula:** 398 M_atp_c + 1990 M_h2o_c + llmg_2195_2mer_degradation_c -> 136 M_ala__L_c + 1990 M_h_c + 398 M_pi_c + 6 M_cys__L_c + 90 M_asp__L_c + 140 M_glu__L_c + 48 M_phe__L_c + 108 M_gly_c + 20 M_his__L_c + 112 M_ile__L_c + 100 M_lys__L_c + 146 M_leu__L_c + 398 M_adp_c + 48 M_gln__L_c + 8 M_trp__L_c + 56 M_arg__L_c + 82 M_asn__L_c + 40 M_tyr__L_c + 92 M_ser__L_c + 44 M_met__L_c + 68 M_pro__L_c + 94 M_thr__L_c + 156 M_val__L_c
- **Catalyst:** Clp_Protease_Complex_Enzyme_c

- **Reaction ID:** R_llmg_2196_2mer_degradation
- **Reaction Formula:** 172 M_atp_c + 860 M_h2o_c + llmg_2196_2mer_degradation_c -> 38 M_ala__L_c + 860 M_h_c + 172 M_pi_c + 8 M_cys__L_c + 48 M_asp__L_c + 62 M_glu__L_c + 34 M_phe__L_c + 58 M_gly_c + 20 M_his__L_c + 36 M_ile__L_c + 44 M_lys__L_c + 66 M_leu__L_c + 172 M_adp_c + 26 M_gln__L_c + 4 M_trp__L_c + 42 M_arg__L_c + 26 M_asn__L_c + 18 M_tyr__L_c + 28 M_ser__L_c + 32 M_met__L_c + 20 M_pro__L_c + 42 M_thr__L_c + 38 M_val__L_c
- **Catalyst:** Clp_Protease_Complex_Enzyme_c

Regarding the special catalyst, 70S ribosome, we described its degradation in this process. The first step is also the breakdown into rRNA and ribosomal proteins:

- **Reaction ID:** R_ribosome_70S_degradation
- **Reaction Formula:** ribosome_70S_c -> rRNA_5S_unmodified_c + rRNA_16S_c + rRNA_23S_c + ribosome_30S_protein_degraded_c + ribosome_50S_protein_degraded_c
- **Catalyst:** -

Then, the second step is the breakdown of 30S ribosomal protein and 50S ribosomal protein into sub-proteins. The resulting rRNA would not be degraded further as they seem to be stable. Below are the reactions:

- **Reaction ID:** R_ribosome_30S_protein_degraded_degradation
- **Reaction Formula:** ribosome_30S_protein_degraded_c -> llmg_0251_Monomer_degradation_c + llmg_0296_Monomer_degradation_c + llmg_0932_Monomer_degradation_c + llmg_1724_assumed_Monomer_degradation_c + llmg_1921_Monomer_degradation_c + llmg_2078_Monomer_degradation_c + llmg_2355_Monomer_degradation_c + llmg_2356_Monomer_degradation_c + llmg_2364_Monomer_degradation_c + llmg_2367_Monomer_degradation_c + llmg_2370_Monomer_degradation_c + llmg_2374_Monomer_degradation_c + llmg_2377_Monomer_degradation_c + llmg_2379_Monomer_degradation_c + llmg_2384_Monomer_degradation_c + llmg_2430_Monomer_degradation_c + llmg_2473_Monomer_degradation_c + llmg_2475_Monomer_degradation_c + llmg_2545_Monomer_degradation_c + llmg_2557_Monomer_degradation_c + llmg_2558_Monomer_degradation_c
- **Catalyst:** -

- **Reaction ID:** R_ribosome_50S_protein_degraded_degradation

- **Reaction Formula:** ribosome_50S_protein_degraded_c -> llmg_0098_Monomer_degradation_c + llmg_0099_Monomer_degradation_c + llmg_0145_Monomer_degradation_c + llmg_0204_Monomer_degradation_c + llmg_0906_Monomer_degradation_c + llmg_1207_Monomer_degradation_c + 4 llmg_1208_Monomer_degradation_c + llmg_1491_Monomer_degradation_c + llmg_1493_Monomer_degradation_c + llmg_1671_Monomer_degradation_c + llmg_1815_Monomer_degradation_c + llmg_2029_Monomer_degradation_c + llmg_2030_Monomer_degradation_c + llmg_2276_Monomer_degradation_c + llmg_2277_Monomer_degradation_c + llmg_2353_Monomer_degradation_c + llmg_2357_Monomer_degradation_c + llmg_2362_Monomer_degradation_c + llmg_2363_Monomer_degradation_c + llmg_2365_Monomer_degradation_c + llmg_2366_Monomer_degradation_c + llmg_2371_Monomer_degradation_c + llmg_2372_Monomer_degradation_c + llmg_2373_Monomer_degradation_c + llmg_2375_Monomer_degradation_c + llmg_2376_Monomer_degradation_c + llmg_2378_Monomer_degradation_c + llmg_2380_Monomer_degradation_c + llmg_2381_Monomer_degradation_c + llmg_2382_Monomer_degradation_c + llmg_2383_Monomer_degradation_c + llmg_2546_Monomer_degradation_c
- **Catalyst:** -

At last, these sub-proteins are degraded into amino acids, which are similar to the proteolysis reactions of other sub-proteins, and examples will thus not be shown here.

#### 3.14 Enzyme dilution

We formulated enzyme dilution reactions in the model to represent the dilution of functional enzymes to daughter cells during cell division. In other words, these reactions partly represent protein composition in a cell. We formulated such reactions for all the catalysts but not for the subunits. Below is an example:

- **Reaction ID:** R_dilution_Clp_Protease_Complex_Enzyme
- **Reaction Formula:** Clp_Protease_Complex_Enzyme_c ->
- **Catalyst:** -

Besides, dilution of 70S ribosome was also included in this process:

- **Reaction ID:** R_dilution_ribosome_70S
- **Reaction Formula:** ribosome_70S_c ->
- **Catalyst:** -

Note that rRNA would not be diluted in the model as its final state 70S ribosome has been diluted.

##### 3.15 RNA dilution

We also formulated RNA dilution reactions in the model to represent the dilution of RNA to daughter cells during cell division. We formulated such reactions only for the TUs used for translation and generic tRNA. Dilution of rRNA is already considered in the dilution of 70S ribosome.

Below is an example of TU dilution:

- **Reaction ID:** R_dilution_mRNA_TU1G1R_100
- **Reaction Formula:** TU1G1R_100_c ->
- **Catalyst:** -

Below is an example of generic tRNA dilution:

- **Reaction ID:** R_dilution_generic_tRNA_Ala_GCA
- **Reaction Formula:** tRNA_Ala_GCA_c ->
- **Catalyst:** -

### 4 Formulation of other reactions

There are some other reactions that should be formulated in the model, including synthesis of unmodelled protein and a biomass dilution reaction with growth-associated maintenance (GAM) involved.

#### 4.1 Adding an unmodelled protein

By considering enzyme and RNA dilution reactions, the model is able to account for the dilution of protein and RNA in biomass. But the model does not contain all the proteins in *L. lactis*. An unmodelled protein should be added to represent the total of the remaining proteins. We assumed that the unmodelled protein had the same amino acid composition as that in biomass equation used in the M model and the number of the amino acids was 250, which is the median length of the proteins in *L. lactis*. Besides, we assumed that the codon of each amino acid was the most frequently used one in *L. lactis* based on the GtRNAdb database[27].

#### 4.2 Adding a biomass dilution reaction

Considering the fact that the model is able to describe the dilution of RNA and proteins to daughter, we formulated the dilution of other biomass materials. The biomass dilution reaction was derived from the biomass equation in the M model, in which protein and RNA were removed as they were represented in the model by dilution reactions. The unmodelled protein was, however, added in the biomass dilution reaction with a specific stoichiometric coefficient. Besides, we assumed that the unmodelled protein accounted for approximately 40% of the *L. lactis* proteome by mass based on published data[19] and then determined the stoichiometric coefficient of the unmodelled protein. We assumed that all these materials in the biomass dilution reaction were growth rate independent.

We calculated GAM and non-growth-associated maintenance (NGAM) in *L. lactis*, as described by[16], using physiological data from different specific growth rates[15]. We calculated that GAM is 34 mmolATP/gCDW and NGAM is 3.7 mmolATP/gCDW/h as can be seen in Fig. 2. Thus we used these two values in the M model.

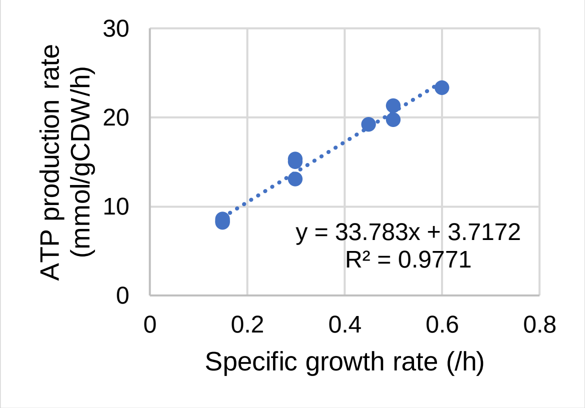

**Fig. 2. Estimation of GAM and NGAM based on experimental data.**

GAM in the proteome-constrained model was calculated by removing the parts contributed by protein synthesis. There are two major ATP consumption processes included in the proteome-constrained model but not in the M model, i.e., tRNA charging and translation. It was estimated that the tRNA charging process accounted for around 3 to 6 mmol ATP per gram CDW, and translation for around 6 to 10 mmol ATP per gram CDW, but the M model itself has already accounted for around 18 mmol ATP per gram CDW. Accordingly, GAM is set as 34 + 18 – 16 = 36 mmol/gCDW. NGAM is decreased to 2 mmol/gCDW/h as energy cost of protein and mRNA turnover have been in part accounted for in the model.

- **Reaction ID:** R_biomass_dilution
- **Reaction Formula:** 36 M_atp_c + 36 M_h2o_c + 0.002 M_nad_c + 0.0002 M_coa_c + 1e-05 M_thf_c + 6.1e-05 M_pg_LLA_c + 0.000138 M_clpn_LLA_c + 0.0064 M_CPS_LLA_c + 9.6e-05 M_d12dg_LLA_c + 0.00074 M_DNA_LLA_c + 0.00015 M_LTAAlaGal_LLA_c + 1.3e-05 M_lyspg_LLA_c + 0.119 M_PG_c + 0.0002 M_udcpdp_c + 1e-05 M_thmpp_c + 1.3e-05 M_m12dg_LLA_c + 0.00650389 unmodelled _protein_biomass_c -> 36 M_h_c + 36 M_pi_c + 36 M_adp_c

- **Catalyst:** -

### 5 Constraints

#### 5.1 Collection of turnover rates for metabolic reactions

Turnover rates, i.e., $k_{cat}$ values, are used for coupling constraints and thus should be collected and assigned. The $k_{cat}$ values of metabolic enzymes were assigned for enzymatic reactions using the GECKO toolbox[28]. Note that the collected $k_{cat}$ values of subunits should be multiplied by the stoichiometry coefficients in enzymes. For the enzymes without assigned $k_{cat}$ values, we assumed them to be the median of the collected ones, which is 100 /s. Besides, we manually assigned a $k_{cat}$ value for glucose transporter based on *E. coli* data[29], which is 180 /s.

#### 5.2 Estimation of catalytic rates for gene expression process

According to a previous study, the equation of ribosomal catalytic rate follows a Michaelis-Menten-type[30], and we assumed that it was also the case in *L. lactis*. Below we show how we obtained parameters and estimated catalytic rates for several machineries.

In *E. coli*, the RNA-to-Protein ratio follows the equation[30]:

$$\frac{R}{P} = \frac{\mu}{\kappa_t} + r_0 \tag{1}$$

in which $R$ is total cellular RNA mass (g/gCDW), $P$ is total cellular protein mass (g/gCDW).

a) $\kappa_t$ and $r_0$ in *L. lactis*

The study[31] showed that there was a linear correlation between specific growth rate and RNA/protein ratio in *L. lactis* NCDO 712 with the slope being 0.0623 while the intercept 0.0342 as shown in Fig. 3. In another study[32] it was found that the total protein accounts for 0.45 g/gCDW and total RNA 0.08 g/gCDW at the growth rate of 0.8 /h. With the relationship in Fig. 3, we estimated total protein and RNA for the growth rate of 0.8 /h. We found that if the total protein is 0.45 g/gCDW, then the total RNA is only 0.0378 g/gCDW, which is lower than the reported value in the study[32]. It seems that the RNA measurements were underestimated at that time. Therefore, we adjusted the relationship with a factor of 0.08/0.0378 = 2.1, i.e., both slope and intercept were increased. As a result, the adjusted slope is 0.13 while intercept 0.072. Accordingly, we can estimate $\kappa_t$ and $r_0$ based on equation (1):

$$\kappa_t = \frac{1}{0.13} = 7.7 \tag{2}$$

$$r_0 = 0.072 \tag{3}$$

Now the equation of ribosomal catalytic rate follows a Michaelis-Menten-type. Therefore, we can calculate the key parameters in the equation if data available for *L. lactis*.

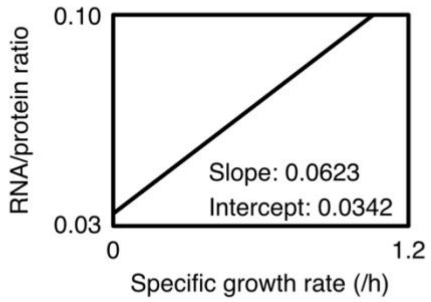

**Fig. 3. Published relationship between specific growth rate and RNA/protein ratio in *L. lactis*.**

b) Ribosomal catalytic rate

Ribosomal catalytic rate (aa/ribosome/s) can be formulated as

$$k_{ribo} = \frac{P_S}{n_r} = \frac{\mu P / m_{aa}}{R f_{rRNA} / m_{rr}} \tag{4}$$

in which $P_S$ is protein synthesis rate (aa/s), $n_r$ is number of ribosomes, $m_{aa}$ is molecular weight of average amino acid (g/mol), $m_{rr}$ is mass of rRNA per ribosome (g/mol_ribosome), $f_{rRNA}$ is fraction of rRNA in total RNA.

Let

$$c_{ribosome} = \frac{m_{rr}}{m_{aa} \cdot f_{rRNA}} \tag{5}$$

then

$$k_{ribo} = \frac{c_{ribosome} \cdot \mu \cdot P}{R} \tag{6}$$

Using the RNA-to-Protein ratio equation, then

$$k_{ribo} = \frac{c_{ribosome} \cdot \kappa_t \cdot \mu}{\mu + r_0 \cdot \kappa_t} = \frac{V_{max} \cdot \mu}{\mu + K_m} \tag{7}$$

in which

$$V_{max} = c_{ribosome} \cdot \kappa_t \tag{8}$$

$$K_m = r_0 \cdot \kappa_t \tag{9}$$

Now the equation of ribosomal catalytic rate follows a Michaelis-Menten-type. Therefore, we can calculate ribosomal catalytic rate for *L. lactis*.

Given that $m_{rr}$ is 1700000 g/mol_ribosome, $m_{aa}$ is 109 g/mol, $f_{rRNA}$ is 0.86, we can calculate the value of $c_{ribosome}$ and then $V_{max}$ and $K_m$:

$$V_{max} = 5.04 \times 7.7 = 38.8 \tag{10}$$

$$K_m = 0.072 \times 7.7 = 0.55 \tag{11}$$

So ribosomal catalytic rate (aa/ribosome/s) is:

$$k_{ribo} = \frac{V_{max} \cdot \mu}{\mu + K_m} = \frac{38.8\mu}{\mu + 0.55} \tag{12}$$

c) RNA polymerase catalytic rate:

In *E. coli*, the transcription rate is exactly 3 times the translation rate at all growth rates[33]. Therefore, the RNA polymerase catalytic rate is three times ribosomal catalytic rate (nucleotide/rnap/s):

$$k_{rnap} = 3k_{ribo} = \frac{116.4\mu}{\mu + 0.55} \tag{13}$$

d) mRNA catalytic rate:

mRNA catalytic rate (protein/mRNA/s) is

$$k_{mRNA} = \frac{c_{mRNA} \cdot \mu \cdot P}{R} = \frac{c_{mRNA} \cdot k_T \cdot \mu}{r_o \cdot k_T + \mu} \tag{14}$$

in which

$$c_{mRNA} = \frac{m_{nt}}{m_{aa} \cdot f_{mRNA}} \tag{15}$$

Given that $m_{nt}$ (molecular weight of average mRNA nucleotide) is 324 g/mol, $m_{aa}$ is 109 g/mol, $f_{mRNA}$ (fraction of mRNA in total RNA) is 0.02, $\kappa_t$ is 7.7 /h, $r_0$ is 0.072, average protein length is 350, average transcription unit length is 1591, we can calculate

$$k_{mRNA} = \frac{1.45\mu}{\mu + 0.55} \tag{16}$$

e) tRNA catalytic rate:

tRNA catalytic rate (aa/tRNA/s) is

$$k_{tRNA} = \frac{c_{tRNA} \cdot \mu \cdot P}{R} = \frac{c_{tRNA} \cdot k_T \cdot \mu}{r_o \cdot k_T + \mu} \tag{17}$$

in which

$$c_{tRNA} = \frac{m_{tRNA}}{m_{aa} \cdot f_{tRNA}} \tag{18}$$

Given that $m_{tRNA}$ (molecular weight of average tRNA) is 25000 g/mol, $m_{aa}$ is 109 g/mol, $f_{tRNA}$ (fraction of tRNA in total RNA) is 0.12 $\kappa_t$ is 7.7 /h, $r_0$ is 0.072, we can calculate

$$k_{tRNA} = \frac{4.1\mu}{\mu + 0.55} \tag{19}$$

#### 5.3 Constraints

Simulations are usually performed by solving linear programming (LP) problems, in which one or several reaction rates are maximized or minimized. The simulation will probably move closer to the real phenotype when more constrains are added. Accordingly, imposing constraints on reaction rates is crucial when performing simulations. We discuss below the constraints used in simulations. In addition to a few basic constraints that are used in the M model, there are some coupling constraints that are imposed to couple different biological processes.

a) Basic constraints

The proteome-constrained model is an expanded version of the M model, and thus is also able to include two basic constraints of the M model. The first one is SV = 0 at the steady-state condition. When performing simulations, we assume that the concentration of each component constant, meaning that the sum of the rates producing it equals the sum of the rates consuming it. The second constraint is the lower and upper bounds on reaction rates. This enables a feasible solution space where the simulated state must be located. For most of the reactions in the model, we lack the data of their bounds, and thereby do not constrain them. However, we can usually constrain a few of them, e.g., exchange reactions, based on external conditions, e.g., medium composition.

These are some common constraints in the M model. Below we discuss some specific constraints used in the proteome-constrained model.

b) Coupling metabolic reactions and enzymes

In reality, any reaction rate in the cell cannot reach the default maximum value in the model, i.e., 1000 mmol/gCDW/h, as the cell cannot maintain enough molecules of the enzyme in the limited intracellular space to hold such a high flux. Therefore, there comes the first coupling constraint, i.e., the rate of a metabolic reaction is constrained by the concentration of the enzyme that catalyses it:

$$V_{met,i} \leq k_{cat,i} \cdot [E]_i \quad (20)$$

in which $V_{met,i}$ represents the rate (mmol/gCDW/h) of the metabolic reaction, $k_{cat,i}$ is the turnover rate (/h), and $[E]_i$ represents the concentration (mmol/gCDW) of the enzyme in the cell. The turnover rate has already been collected and can thus be used directly. Below we focus on the enzyme concentration. At the steady-state condition, the concentration of the enzyme is constant:

$$d[E]_i/dt = V_{syn,i} - V_{deg,i} - V_{dil,i} = 0 \quad (21)$$

in which $V_{syn,i}$ represents the synthesis rate (mmol/gCDW/h) of the enzyme, $V_{deg,i}$ represents its degradation rate (mmol/gCDW/h), and $V_{dil,i}$ represents its dilution rate (mmol/gCDW/h). Equation (21) can then be converted to

$$V_{syn,i} - k_{deg,enzyme,i} \cdot [E]_i - \mu \cdot [E]_i = 0 \quad (22)$$

$$[E]_i = \frac{1}{k_{deg,enzyme,i} + \mu} \cdot V_{syn,i} \quad (23)$$

in which, $k_{deg,i}$ is degradation constant (/h), and $\mu$ is growth rate (/h). By combining (20) and (23), we have

$$V_{met,i} \leq \frac{k_{cat,i}}{k_{deg,enzyme,i} + \mu} \cdot V_{syn,i} \tag{24}$$

Accordingly, inequation (25) is used for coupling the metabolic reactions and enzyme formation reactions.

For the case that one enzyme catalyses two or more metabolic reactions, inequation can also be generated. For example, if an enzyme can catalyse two metabolic reactions, we can have

$$\frac{1}{k_{cat,1}} \cdot V_{met,1} + \frac{1}{k_{cat,2}} \cdot V_{met,2} \leq \frac{1}{k_{deg,enzyme} + \mu} \cdot V_{syn} \tag{25}$$

There is, however, no such case in the metabolism part as we assigned the man-made enzyme, based on the reaction ID, for each enzymatic reaction in the M model.

Note that there will be some cases that optimal solution cannot be achieved probably due to some extremely low $k_{cat}$ values, we assume the $k_{cat}$ values below 10% lowest $k_{cat}$ values (5551 /h) to be 5551 /h.

c) Coupling some gene expression reactions and enzymes

Similar to metabolic reactions, some reactions in gene expression processes can also be coupled with the enzyme synthesis reactions using the turnover rate as one of the coupling parameters. These enzymes include tRNA synthetases, methionine aminopeptidase, peptide deformylase, and enzymes in RNA modification process. These reactions follow the same way as metabolic reactions to be coupled with their enzyme formation reactions. We used the median $k_{cat}$ value in the metabolic part for the protein synthesis part. Moreover, the catalytic rates of some catalysts depend on the length of the substrates (e.g., the catalytic rate of RNA polymerase relies on the length of the TU and thereby differs from TUs with various sequences) and growth rate. Next, we will show coupling constraints about the specific machineries.

d) Coupling transcription reactions and RNA polymerase

In the model, we use the RNA polymerase together with the sigma factor as the complex to catalyse transcription reactions. The catalytic rate (nucleotide molecules/complex molecule/s) of RNA polymerase changes with growth rate as shown in equation (13). Given that the catalytic rate is for nucleotides, we can have the catalytic rate for TU molecules, which should be divided by the number of nucleotides in the TU:

$$k_{rnap\&\sigma,i} = \frac{\frac{176.46 \cdot \mu}{1.147 + \mu}}{length(TU_i)} \tag{26}$$

which has the unit of "TU molecules/complex molecule/s". By balancing the concentration of the complex of RNA polymerase with sigma factor, we can have:

$$\sum \frac{V_{transcription\ of\ TU,i}}{k_{rnap\&\sigma,i}} \le \frac{1}{k_{deg,rnap\&\sigma} + \mu} \cdot V_{syn,rnap\&\sigma} \tag{27}$$

Then we can have:

$$\sum (V_{transcription\ of\ TU,i} \cdot length(TU_i)) \le \frac{k_{rnap\&\sigma}}{k_{deg,rnap\&\sigma} + \mu} \cdot V_{syn,rnap\&\sigma} \tag{28}$$

e) Coupling mRNA degradation reactions and mRNA degradation complex

In the model, the mRNA degradation complex is made up of degradosome together with oligoribonuclease. Similarly, we can have the constraint:

$$\sum (V_{degradation\ of\ TU,i} \cdot length(TU_i)) \le \frac{k_{mrnadegcplx}}{k_{deg,mrnadegcplx} + \mu} \cdot V_{syn,mrnadegcplx} \tag{29}$$

We assume that the catalytic rate of the mRNA degradation complex is calculated as follow:

$$k_{mrnadegcplx} = 100 \times 1591 \tag{30}$$

in which the median $k_{cat}$ value of metabolic enzymes is 100 /s, average transcription unit length is 1591. The unit is nucleotide molecules/complex molecule/s.

f) Coupling tRNA

As we have not added tRNA as reactants in the tRNA charging and translation reactions, tRNA production reactions, i.e., generic RNA renaming reactions, and tRNA charging reactions should be coupled to enable active fluxes of tRNA. tRNA can be regarded as the catalyst in tRNA charging reactions, and thereby assigned with the catalytic rate, which has been shown in equation (19). Accordingly, tRNA charging reaction is coupled with not only the synthesis of tRNA synthetase, but also the production of tRNA. Below show the two coupling constraints:

$$V_{trnacharging,i} \le \frac{k_{cat,synthetase,i}}{k_{deg,synthetase,i} + \mu} \cdot V_{syn,synthetase,i} \tag{31}$$

$$V_{trnacharging} \le \frac{k_{trna}}{\mu} \cdot \sum V_{trnarenaming,i} \tag{32}$$

Note that tRNA is assumed to be stable and thereby be not degraded. So, there is no degradation factor in inequation (32).

g) Coupling mRNA

There are two types of TUs in the model, one is for translation, which will be degraded and diluted, and the other is for stable RNA cleavage, which will be cleaved.

Regarding the TUs for translation, we have not added them as reactants in the translation reactions, so TU production and translation reactions should be coupled. The catalytic rate of mRNA has been shown in equation (16). Theoretically, TUs can be produced by transcription reactions and stable RNA cleavage reactions. There is, however, no TU from stable RNA cleavage reactions that contain intact genes in the model. So, we can just couple translation initiation reactions and transcription reactions:

$$\sum V_{translationinitiation,i} \leq \frac{k_{mrna}}{k_{deg,mrna,i} + \mu} \cdot V_{transcription} \quad (33)$$

Regarding the TUs for stable RNA cleavage, transcription rate equals to cleavage rate at the steady-state condition. There is neither degradation nor dilution reactions for them.

h) Coupling translation reactions and ribosome

In addition to mRNA, translation reactions are constrained by ribosome production. Therefore, translation reaction should also be coupled with ribosome production reaction. We couple the translation elongation reactions with ribosome. The catalytic rate of 70S ribosome has been shown in equation (12). Similar to the RNA polymerase case, the coupling constraint can be set as:

$$\sum (V_{translationelongation,i} \cdot length(peptide_i)) \leq \frac{k_{ribo}}{k_{deg,ribo} + \mu} \cdot V_{syn,ribo} \quad (34)$$

i) Coupling protein degradation reactions and protease

We assume that the catalytic rate of the protease complex is calculated as follow:

$$k_{protease} = 100 \times 350 \quad (35)$$

in which the median $k_{cat}$ value of metabolic enzymes is 100 /s, average protein length is 350. The unit is amino acid molecules/complex molecule/s. Similar to mRNA degradation complex, we can have the constraint:

$$\sum (V_{degradation\ of\ protein,i} \cdot length(protein_i)) \leq \frac{k_{protease}}{k_{deg,protease} + \mu} \cdot V_{syn,protease} \quad (36)$$

j) Degradation of mRNA, 70S ribosome and enzymes

The model accounts for degradation of mRNA, 70S ribosome and enzymes. All these degradation reactions are assumed to follow the first order rate constant:

$$V_{deg,mrna,i} = k_{deg,mrna,i} \cdot [TU]_i = k_{deg,mrna,i} \cdot \frac{1}{k_{deg,mrna,i} + \mu} \cdot V_{transcription,i} \quad (37)$$

$$V_{deg,ribo} = k_{deg,ribo} \cdot [Ribo]_i = k_{deg,ribo} \cdot \frac{1}{k_{deg,ribo} + \mu} \cdot V_{syn,ribo} \quad (38)$$

$$V_{deg,enzyme,i} = k_{deg,enzyme,i} \cdot [E]_i = k_{deg,enzyme,i} \cdot \frac{1}{k_{deg,enzyme,i} + \mu} \cdot V_{syn,enzyme,i} \quad (39)$$

For mRNA degradation, the median half-lives of mRNA in *L. lactis* were measured to be 5.8, 11.4 and 15.5 min, at 0.80, 0.51 and 0.11 /h, respectively[34]. Accordingly, the degradation constant of mRNA is:

$$k_{deg,mrna} = 6.25 \times \mu + 1.54 \tag{40}$$

In the model, we assume the global protein degradation constant (1/h) to be:

$$k_{deg,enzyme} = 0.01\mu \tag{41}$$

and ribosome degradation constant to be:

$$k_{deg,ribo} = 0.01\mu \tag{42}$$

Note that the TUs used for translation should be degraded, TUs for cleavage should not be degraded, and sub-TUs should also be degraded. Besides, the unmodelled protein should be degraded. The coupling constraint follows:

$$V_{deg,ump} = k_{deg,ump} \cdot \frac{1}{k_{deg,ump} + \mu} \cdot V_{syn,ump} \tag{43}$$

k) Dilution of mRNA, ribosome and enzymes

The model also accounts for the dilution of mRNA, tRNA, ribosome and enzymes into daughter cells. The dilution constant rate is the growth rate. Then the dilution rates of them should be:

$$V_{dil,mrna,i} = \mu \cdot [TU]_i = \mu \cdot \frac{1}{k_{deg,mrna,i} + \mu} \cdot V_{transcription,i} \tag{44}$$

$$V_{dil,ribo} = \mu \cdot [Ribo] = \mu \cdot \frac{1}{k_{deg,ribo} + \mu} \cdot V_{syn,ribo} \tag{45}$$

$$V_{dil,enzyme,i} = \mu \cdot [E]_i = \mu \cdot \frac{1}{k_{deg,enzyme,i} + \mu} \cdot V_{syn,enzyme,i} \tag{46}$$

$$V_{dil,ump} = \mu \cdot \frac{1}{k_{deg,ump} + \mu} \cdot V_{syn,ump} \tag{47}$$

As there is no reaction for tRNA degradation in the model, all the tRNA will be diluted. So there is no need to formulate coupling. Besides, the TUs used for translation should be diluted, TUs for cleavage should not, but sub-TUs should also be diluted.

l) Constraint of total protein and RNA

There is a biomass dilution reaction representing a constant biomass composition, which means that most of the components, except protein and RNA, are assumed to be growth rate independent. On the other hand, the total protein pool is assumed to be fixed at 0.46 g/gCDW, i.e.,

$$\sum [E]_i + [Ribosomal\, proteins] = 0.46 \tag{48}$$

Besides, we do not constrain the total RNA pool in the model, which is determined by the total protein requirement together with catalytic rates of ribosome, tRNA and mRNA for each specific simulation.

m) Constraint of glucose transporter

Due to the limited space of membrane or some regulatory mechanisms, the glucose transporter also has an upper limit. We assume in the model that the total concentration of glucose transporter should be not greater than a given value which can be estimated by the model or from literatures, i.e.,

$$[Glucose\ transporter] \leq N \tag{49}$$

$$\frac{V_{dil,glucose\ transporter}}{\mu} \leq N \tag{50}$$

$$V_{dil,glucose\ transporter} \leq \mu \cdot N \tag{51}$$

**Reference:**

1. Heirendt, L. *et al.* Creation and analysis of biochemical constraint-based models using the COBRA Toolbox v.3.0. *Nat. Protoc.* 1 (2019).
2. O'Brien, E. J., Lerman, J. A., Chang, R. L., Hyduke, D. R. & Palsson, B. Genome-scale models of metabolism and gene expression extend and refine growth phenotype prediction. *Mol. Syst. Biol.* **9**, 693 (2013).
3. Lerman, J. A. *et al.* In silico method for modelling metabolism and gene product expression at genome scale. *Nat. Commun.* **3**, 929 (2012).
4. Huerta-Cepas, J. *et al.* eggNOG 4.5: a hierarchical orthology framework with improved functional annotations for eukaryotic, prokaryotic and viral sequences. *Nucleic Acids Res.* **44**, (2016).
5. Karp, P. D. *et al.* The BioCyc collection of microbial genomes and metabolic pathways. *Brief. Bioinform.* **20**, 1085–1093 (2019).
6. Puri, P. *et al.* Systematic identification of tRNAome and its dynamics in Lactococcus lactis. *Mol. Microbiol.* **93**, 944–956 (2014).
7. Machnicka, M. A., Dunin-Horkawicz, S., de Crécy-Lagard, V. & Bujnicki, J. M. tRNAmodpred: A computational method for predicting posttranscriptional modifications in tRNAs. *Methods* **107**, 34–41 (2016).
8. Larkin, M. A. *et al.* Clustal W and Clustal X version 2.0. **23**, 2947–2948 (2007).
9. Thiele, I., Jamshidi, N., Fleming, R. M. T. & Palsson, B. Ø. Genome-Scale Reconstruction of Escherichia coli's Transcriptional and Translational Machinery: A Knowledge Base, Its Mathematical Formulation, and Its Functional Characterization. *PLoS Comput. Biol.* **5**, e1000312 (2009).

10. Burley, S. K. *et al.* Protein Data Bank: the single global archive for 3D macromolecular structure data. *Nucleic Acids Res.* **47**, D520–D528 (2019).
11. UniProt Consortium, T. UniProt: the universal protein knowledgebase. *Nucleic Acids Res.* **46**, 2699–2699 (2018).
12. Kanehisa, M., Sato, Y., Furumichi, M., Morishima, K. & Tanabe, M. New approach for understanding genome variations in KEGG. *Nucleic Acids Res.* **47**, (2019).
13. Martinez, A. *et al.* Extent of N-terminal modifications in cytosolic proteins from eukaryotes. *Proteomics* **8**, 2809–2831 (2008).
14. Flahaut, N. A. L. *et al.* Genome-scale metabolic model for Lactococcus lactis MG1363 and its application to the analysis of flavor formation. *Appl. Microbiol. Biotechnol.* **97**, 8729–8739 (2013).
15. Goel, A. *et al.* Protein costs do not explain evolution of metabolic strategies and regulation of ribosomal content: does protein investment explain an anaerobic bacterial Crabtree effect? *Mol. Microbiol.* **97**, 77–92 (2015).
16. Teusink, B. *et al.* Analysis of growth of Lactobacillus plantarum WCFS1 on a complex medium using a genome-scale metabolic model. *J. Biol. Chem.* **281**, 40041–8 (2006).
17. King, Z. A. *et al.* BiGG Models: A platform for integrating, standardizing and sharing genome-scale models. *Nucleic Acids Res.* **44**, 515–522 (2015).
18. Mendoza, S. N., Olivier, B. G., Molenaar, D. & Teusink, B. A systematic assessment of current genome-scale metabolic reconstruction tools. *Genome Biol.* **20**, 158 (2019).
19. Lahtvee, P.-J., Seiman, A., Arike, L., Adamberg, K. & Vilu, R. Protein turnover forms one of the highest maintenance costs in Lactococcus lactis. *Microbiology* **160**, 1501–1512 (2014).
20. Mulder, F. A. A. *et al.* Conformation and dynamics of ribosomal stalk protein L12 in solution and on the ribosome. *Biochemistry* **43**, 5930–5936 (2004).
21. Frees, D. & Ingmer, H. ClpP participates in the degradation of misfolded protein in Lactococcus lactis. *Mol. Microbiol.* **31**, 79–87 (1999).
22. Nilsson, D., Lauridsen, A. A., Tomoyasu, T. & Ogura, T. A Lactococcus lactis gene encodes a membrane protein with putative ATPase activity that is homologous to the essential Escherichia coli ftsH gene product. *Microbiology* **140**, 2601–2610 (1994).
23. Poquet, I. *et al.* HtrA is the unique surface housekeeping protease in Lactococcus lactis and is required for natural protein processing. *Mol. Microbiol.* **35**, 1042–1051 (2000).

24. Savijoki, K., Ingmer, H. & Varmanen, P. Proteolytic systems of lactic acid bacteria. *Appl. Microbiol. Biotechnol.* **71**, 394–406 (2006).
25. Varmanen, P., Ingmer, H. & Vogensen, F. K. ctsR of Lactococcus lactis encodes a negative regulator of clp gene expression. *Microbiology* **146**, 1447–1455 (2000).
26. Lynch, M. & Marinov, G. K. The bioenergetic costs of a gene. *Proc. Natl. Acad. Sci. U. S. A.* **112**, 15690–5 (2015).
27. Chan, P. P. & Lowe, T. M. GtRNAdb 2.0: an expanded database of transfer RNA genes identified in complete and draft genomes. *Nucleic Acids Res.* **44**, (2016).
28. Sánchez, B. J. *et al.* Improving the phenotype predictions of a yeast genome-scale metabolic model by incorporating enzymatic constraints. *Mol. Syst. Biol.* **13**, 935 (2017).
29. Szenk, M., Dill, K. A. & de Graff, A. M. R. Why Do Fast-Growing Bacteria Enter Overflow Metabolism? Testing the Membrane Real Estate Hypothesis. *Cell Syst.* **5**, 95–104 (2017).
30. Scott, M., Gunderson, C. W., Mateescu, E. M., Zhang, Z. & Hwa, T. Interdependence of cell growth and gene expression: origins and consequences. *Science* **330**, 1099–102 (2010).
31. Beresford, T. & Condon, S. Physiological and genetic regulation of rRNA synthesis in Lactococcus. *J. Gen. Microbiol.* **139**, 2009–2017 (1993).
32. Novák, L. & Loubiere, P. The metabolic network of Lactococcus lactis: Distribution of 14C- labeled substrates between catabolic and anabolic pathways. *J. Bacteriol.* **182**, 1136–1143 (2000).
33. Proshkin, S., Rachid Rahmouni, A., Mironov, A. & Nudler, E. Cooperation between translating ribosomes and RNA polymerase in transcription elongation. *Science (80-. ).* **328**, 504–508 (2010).
34. Dressaire, C. *et al.* Role of mRNA Stability during Bacterial Adaptation. *PLoS One* **8**, e59059 (2013).
